# Cancer mutations rewire the RNA methylation specificity of METTL3-METTL14

**DOI:** 10.1101/2023.03.16.532618

**Authors:** Chi Zhang, Luiza Tunes, Meng-Hsiung Hsieh, Ping Wang, Ashwani Kumar, Brijesh B. Khadgi, YenYu Yang, Katelyn A. Doxtader, Emily Herrell, Oliwia Koczy, Rohit Setlem, Xunzhi Zhang, Bret Evers, Yinsheng Wang, Chao Xing, Hao Zhu, Yunsun Nam

**Affiliations:** Department of Biochemistry, Department of Biophysics, Simmons Comprehensive Cancer Center, University of Texas Southwestern Medical Center, Dallas, TX 75390, USA; Children’s Research Institute, Departments of Pediatrics and Internal Medicine, Center for Regenerative Science and Medicine, University of Texas Southwestern Medical Center, Dallas, TX 75390, USA; Eugene McDermott Center for Human Growth and Development, Department of Bioinformatics, University of Texas Southwestern Medical Center, Dallas, TX 75390, USA; Department of Chemistry, University of California at Riverside, Riverside, CA 92521, USA; Cecil H. and Ida Green Center for Reproductive Biology Sciences, University of Texas Southwestern Medical Center, Dallas, TX 75390, USA; Department of Pathology, University of Texas Southwestern Medical Center, Dallas, TX 75390, USA

## Abstract

Chemical modification of RNAs is important for post-transcriptional gene regulation. The METTL3-METTL14 complex generates most *N*^6^-methyladenosine (m^6^A) modifications in mRNAs, and dysregulated methyltransferase expression has been linked to numerous cancers. Here we show that changes in m^6^A modification location can impact oncogenesis. A gain-of-function missense mutation found in cancer patients, METTL14^R298P^, promotes malignant cell growth in culture and in transgenic mice. The mutant methyltransferase preferentially modifies noncanonical sites containing a GGAU motif and transforms gene expression without increasing global m^6^A levels in mRNAs. The altered substrate specificity is intrinsic to METTL3-METTL14, helping us to propose a structural model for how the METTL3-METTL14 complex selects the cognate RNA sequences for modification. Together, our work highlights that sequence-specific m^6^A deposition is important for proper function of the modification and that noncanonical methylation events can impact aberrant gene expression and oncogenesis.

## INTRODUCTION

Specific and controlled modification of mRNAs with *N*^6^-methyladenosine (m^6^A) is essential for proper gene expression (1, 2). METTL3 and METTL14 form a heterodimer that acts as the prevalent methyltransferase responsible for most m^6^A modifications in mRNAs (3–5), and aberrant expression of METTL3-METTL14 can directly impact mRNA m^6^A modification levels (3, 6). Many cancers have been linked to METTL3-METTL14 overexpression and increased global m^6^A levels in mRNAs, but the underlying molecular mechanism is unclear (7–9). Transcriptome-wide mapping of m^6^A sites affected by METTL3-METTL14 has shown that the preferred RNA targets of METTL3-METTL14 contain the motif DRACH (D, G/A/U; R, G/A; H, A/U/C), with a dominant Cyt following the methylated Ade (10–13). However, the molecular basis for the observed RNA target sequence preference of the major m^6^A writer complex is also lacking. Furthermore, the biological consequence of changing the substrate preference is unknown.

In METTL14, R298 is the most frequently mutated residue in cancer samples (Catalogue of Somatic Mutations in Cancer, COSMIC) (14) (fig. S1, A and B). METTL3-METTL14 structures exhibit a prominent basic patch in METTL14 that may be used to bind RNAs, and R298 is near the center of the potential RNA binding surface (4, 5, 15). However, how the cognate RNA sequences are recognized is unknown. Other than overexpression of the methyltransferase, a way to produce unwanted methylation is to reprogram the methyltransferase to add m^6^A to novel targets. Such novel functions caused by gain-of-function (GOF) mutations can be difficult to detect, although they potentially drive pathogenesis. For example, R298 may be presumed to be a loss-of-function mutation due to its reduced methylation activity on canonical RNA targets (4, 7). Because the core mechanisms underlying substrate specificity of key RNA modification enzymes remain unknown, it is difficult to understand how RNA modification changes contribute to disease.

## RESULTS

### METTL14^R298P^ causes more malignant cell growth in culture and in mice

The frequency of mutations at R298 of METTL14 led us to hypothesize that R298 mutations may have GOF effects that promote oncogenesis. To test the difference between wild-type and mutant METTL14, we generated four stable HepG2 liver cancer cell lines by integrating lentivirus cassettes expressing EGFP, METTL14^WT^, METTL14^R298P^, or METTL14^D312A^ (fig. S1C). D312A is spatially close to R298P but is not derived from cancer patients, and both mutants have lower methylation activity than the wild-type enzyme, using canonical substrates containing the “GGACU” motif (4). A null background or homozygous mutation of the METTL14 gene was not possible to establish despite multiple attempts with genome-editing techniques because of cell viability. Nevertheless, overexpressing mutant METTL14 alleles as EGFP fusion constructs allowed us to assess their role in a meaningful way because METTL14 and METTL3 need to heterodimerize for enzymatic activity. Purified EGFP fusion constructs have similar methylation activity as the isolated wild-type methyltransferase in quantitative enzymatic assays (fig. S1D). None of the stable cell lines exhibited increased global m^6^A levels, likely because of the limited amount of endogenous METTL3 (fig. S1E). In fact, for cells expressing mutant METTL14, m^6^A levels of mRNAs decreased, which might seem like a loss-of-function (LOF) phenotype in the absence of additional information (fig. S1F) (7).

To test the effect of the R298 mutation on cell growth, we tested proliferation, migration, and invasion of the four stable HepG2 cell lines expressing different METTL14 alleles. While the cell proliferation rate was unaffected by the mutations (fig. S1G), cells expressing METTL14^R298P^ showed increased capacity to migrate and invade, whereas cells expressing METTL14^D312A^ showed similar properties as the ones expressing the wild-type protein (Fig. 1, A and B). To test how the changes in cell migration and invasion might affect tumorigenesis in animals, we used hydrodynamic transfection (HDT) to genomically integrate transposons containing METTL3 and METTL14 into mouse livers. To detect differences between wild-type and mutant METTL14 constructs, we used mice with a unique genetic background (liver-specific *Albumin-Cre; Tp53^Fl/Fl^; Lin28a^Fl/Fl^; Lin28b^Fl/Fl^)* to tune tumorigenesis (Fig. 1C). Overexpression of METTL3 and METTL14^R298P^ resulted in larger and more frequent tumors than those injected with METTL3 and METTL14^WT^ (Fig. 1, D and E). Tumors in the METTL14^R298P^-injected group affected a greater portion of the liver, where much of the normal tissue was replaced. The tumors in mice with METTL14^R298P^ also caused a ~4-fold increase in liver mass when compared to the animals injected with the wild-type or D312A variants of METTL14. Histological examination revealed multiple tumors with sarcomatoid features, including atypical, spindly tumor cells with pleomorphic nuclei and increased mitoses (Fig. 1F). Compared to tumors associated with METTL14^WT^ or METTL14^D312A^, METTL14^R298P^ tumors were larger and replaced much of the hepatic parenchyma. Additionally, the sarcomatoid tumors from the mice injected with METTL14^R298P^ showed areas of tumor necrosis, indicative of increased tumor aggressiveness. Therefore, the R298P mutation of METTL14 drives a more malignant growth phenotype than the wild-type protein in cultured cells and in mice.

**Fig. 1.**
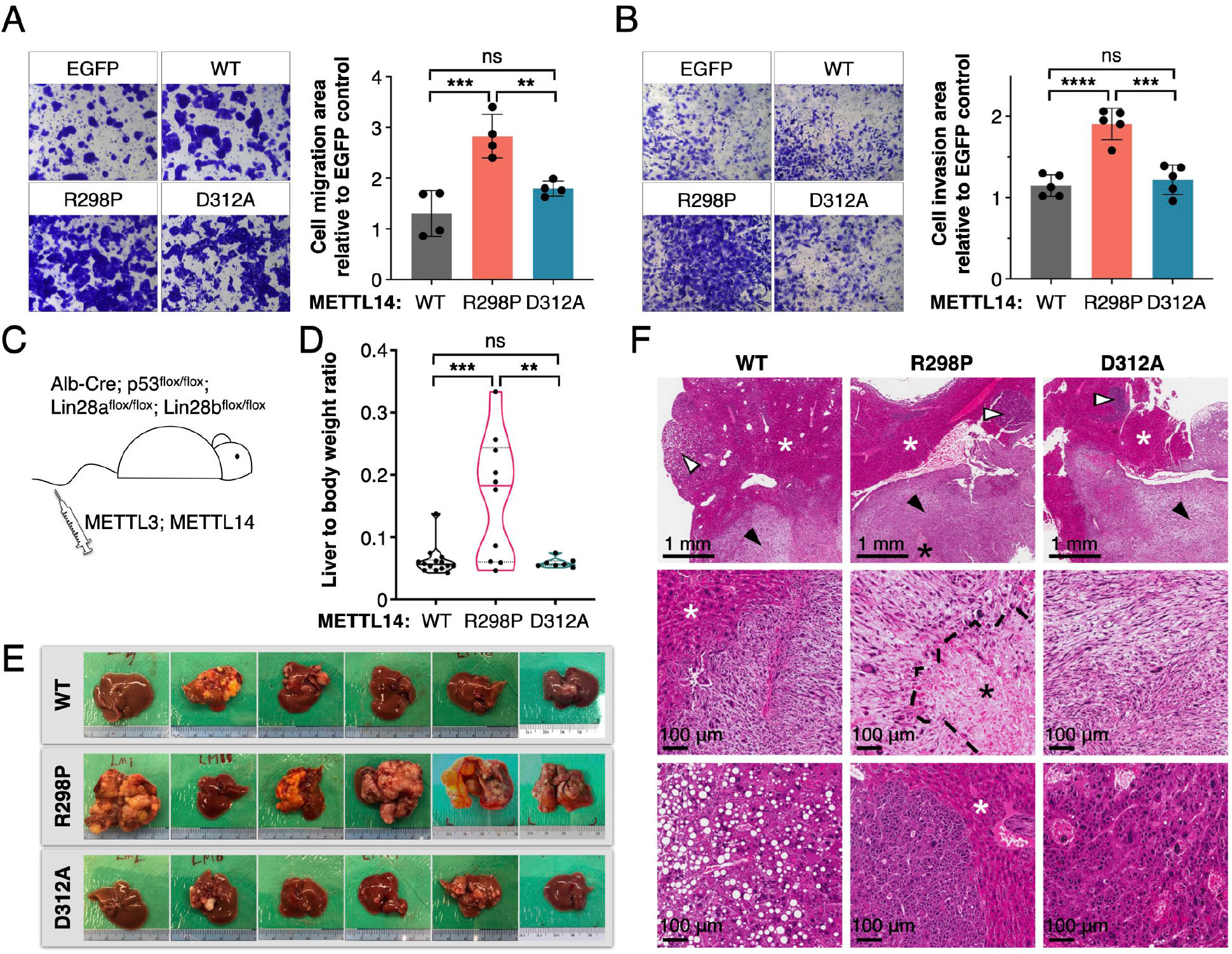
METTL14^R298P^ causes more malignant cell growth in culture and in mice. (**A**and **B**) cell migration assay (A) and invasion assay (B) using HepG2 stable cell lines expressing EGFP or METTL14 (WT, R298P or D312A). The representative bright-field microscopy pictures show the cells stained with crystal violet, and the bar graphs show the migration area or invasion area normalized to the EGFP control. Data are represented as mean ± s.d. from 4(A) or 5(B) biological replicates. Significant difference among means is found in ordinary one-way ANOVA test. Adjusted P values are as indicated: ns, not significant; **, *P* < 0.01; ***, *P* < 0.001; *P* < 0.0001 (Tukey’s multiple comparison test). **(C)** Schematic of the mouse tumorigenesis assay via hydrodynamic transfection. **(D)** Truncated violin plot of the liver-to-body weight ratio of mice 42 days after injection. Data are represented as median and quartiles. The experiment was repeated by using 2-3 cohorts at different times. WT, n=14; R298P, n=10 with two deceased on 35- and 38-day postinjection; D312A, n=7. Adjusted P values from ordinary one-way ANOVA test are as indicated: ns, not significant; **, *P* = 0.0025; ***, *P* = 0.0005 (Tukey’s multiple comparison test). (E) Representative whole liver and tumor tissue images. The replicate numbers are the same as (D). **(F)** Photomicrographs of mouse livers after the tumorigenesis assay. **Upper panels**, large tumors with sarcomatoid features (black arrowheads) replace much of the uninvolved hepatic parenchyma (white asterisks) in R298P mutants compared to D312A mutants and WT controls. Areas of necrosis (black asterisk) are seen within the large sarcomatoid tumors in R298P mutants. Smaller, epithelioid tumors (white arrowheads) are seen in all three groups. **Middle panels**, higher magnification photomicrographs of the sarcomatoid tumors in all three groups reveal atypical, spindly cells with pleomorphic nuclei. Necrosis (black asterisk; dashed line indicates border) is prominent in the tumors from the R298P mutants. **Lower panels**, higher magnification photomicrographs of the epithelioid tumors in all three groups show atypical cells with abundant cytoplasm and pleomorphic nuclei. Scale bars equivalent to 1 mm (upper panel) or 100 μm (middle and lower panels) are shown as black bars.

To understand the oncogenic phenotype, we used RNA-seq analysis to examine the gene expression changes unique to the R298P cell line. Gene Ontology (GO) analysis revealed that certain pathways are consistently differentially expressed in all three pairwise comparisons with R298P. Most notably, the WNT signaling pathway is inhibited in R298P cells compared to the other three cell lines, and multiple groups of genes involved in cell movement or invasion are upregulated (fig. S2, A and B). We confirmed differential expression of certain genes in these top GO term groups using quantitative RT-PCR (fig. S2, C and D). The proteins differentially expressed in cells expressing METTL14^R298P^ compared to the other cell lines can be mapped to similar top GO categories as in the RNA-seq analysis, despite the lower coverage (fig. S2E). Due to the ability of METTL14^R298P^ expression to cause unique gene expression and aggressive growth phenotypes when compared to wild type or another LOF mutant, we surmised that R298 mutations may furnish the methyltransferase with a novel function.

### METTL14^R298P^ exhibits a distinct RNA sequence preference

The R298 residue of METTL14 is located in the middle of a large basic patch that is likely to interact with RNA. Thus, we postulated that mutating R298 might alter RNA substrate specificity. To determine the substrate specificity in an unbiased fashion, we developed the *in-vitro*-methylation-sequencing (IVM-seq) workflow by adopting the strategies from the Systematic Evolution of Ligands by Exponential Enrichment (SELEX) approach (Fig. 2A) (16). IVM-seq allows us to perform a deep search of the sequence space because the complexity of the initial randomized RNA pool (approximately 1×10^12^) is higher than that of a typical transcriptome. In contrast to SELEX which relies on protein-RNA binding affinity to enrich the cognate RNA sequence, we first allowed the methyltransferase to modify target RNAs and then selected the methylated species using an m^6^A antibody. The high-throughput sequencing results were then analyzed to derive the consensus motif (Fig. 2B). The preferred sequence of the wild-type methyltransferase, GGAC, resembled the canonical DRACH motif (10, 11). In contrast, a distinct consensus sequence—GGAU—was derived for METTL3-METTL14^R298P^. IVM-seq analysis of other R298 mutants (R298C and R298H, both found in patients) yielded similar motifs where Ura is preferred over Cyt at the 4th position. Thus, a single amino acid substitution was sufficient to dramatically alter the preferred nucleobase at the 4th position, from Cyt to Ura, changing the consensus motif from GGAC to GGAU. METTL3-METTL14^D312A^ maintained a clear preference for Cyt over Ura at the 4th position, even though the overall low methylation level made it difficult to determine the sequence preference at the first position. Thus, METTL14^D312A^ follows the pattern of a LOF mutation that causes lower methylation for both normal and other targets, whereas the GOF R298 mutations gain increased methylation activity with GGAU targets. To test for potential bias introduced by using a particular antibody, we used multiple anti-m^6^A antibodies produced by different companies (SySy, 202003 and Abcam, ab151230), but the antibody source did not change the sequence preference derived from IVM-seq (Fig. 2C). Using the same randomized RNA library, we also performed SELEX. Unlike IVM-seq, we could not identify a clear consensus sequence for the best binders of METTL3-METTL14 (Fig. 2D). The changes in sequence preference may be more relevant for catalysis than for binding. Therefore, we established robust methods to determine the sequence preference of an RNA m^6^A methyltransferase in an unbiased manner, and we determined that R298 substitutions transform the intrinsic substrate specificity of METTL3-METTL14 independent of other factors.

**Fig. 2.**
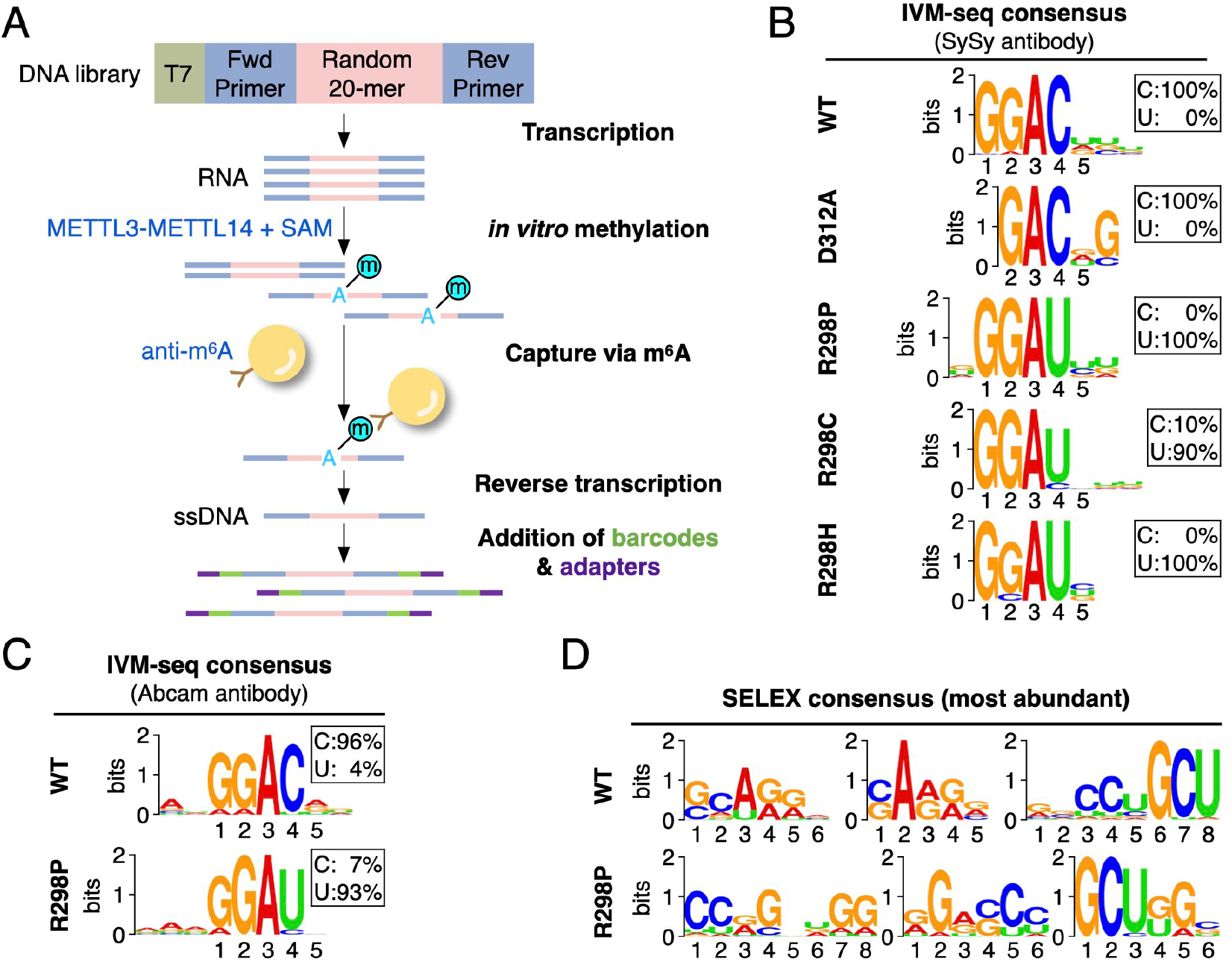
METTL14^R298P^ exhibits a distinct RNA sequence preference. **(A)** Workflow for *in vitro* methylation-sequencing (IVM-seq). Oligonucleotides with a 20-nt randomized segment are used to generate the RNA pool for *in vitro* methylation. RNAs selected with an anti-m^6^A antibody are used for library generation and high-throughput sequencing. (B) Motif analysis of the IVM-seq output for RNAs methylated by the METTL3-METTL14 complexes containing the indicated METTL14, using SySy (202003) anti-m^6^A antibody. The probability percentages of Cyt and Ura at the 4^th^ position are shown in each box. MEME P values are shown top to bottom: 5.2e^-412^, 4.6e^-7^, 4.0e^-355^, 6.5e^-1771^, 6.9e^-1094^. **(C)** Consensus sequence analysis of the IVM-seq output for RNAs methylated by METTL3-METTL14 using Abcam (ab151230) anti-m^6^A antibody. The probability percentages of Cyt and Ura at the 4^th^ position are shown in each box. The Pvalues of the consensus analyzed by MEME are: 1.5e^-32269^ (WT), 9.1e^-45993^ (R298P). (D) The affinity binding consensus sequence derived from the SELEX experiment performed with recombinant METTL3-METTL14 (wild-type or R298P). The P values of the consensus analyzed by MEME are: (top, from left to right) 3.7e^-1045^, 1.5e^-894^,1.3e^-8826^, (bottom, from left to right) 2.0e^-899^, 1.1e^-5651^, 2.5e^-10385^.

To investigate the altered RNA specificity of the METTL3-METTL14 methyltransferase using an orthogonal approach, we used a quantitative enzymatic assay where an established canonical m^6^A site sequence in the *MALAT1* ncRNA was changed combinatorially (Fig. 3A) (17). For the recombinant methyltransferase, we tested patient-derived mutants and additional substitutions that can be obtained through a single nucleotide change. The wild-type enzyme consistently shows a strong preference for Cyt at the 4th position, although some methylation activity is observed for substrates with Ura in the 4th position. In contrast, the substrate preference was reversed for METTL3-METTL14^R298X^ mutants because GGAU substrates were methylated more efficiently than GGAC substrates. Remarkably, the mutant enzyme activity on GGAU substrates is comparable to the wild-type enzyme on the normal sites containing GGAC as indicated by the raw DPM counts. The 5th position is not as discriminatory, but pyrimidines are generally preferred by both wild-type and mutant enzymes. For methyltransferase complexes containing METTL14^WT^ or METTL14^D312A^, the preference for GGAC over GGAU is 3.7- or 6.3-fold, respectively (Fig. 3B). In contrast, all R298 mutations of METTL14 cause a similar change in the preferred substrate sequence, where having Ura at the fourth position increases methylation by 2.4- to 3.6-fold compared with Cyt. While the sequence seems important, the structural context can also affect RNA methylation efficiency (18). Thus, we tested three other RNA scaffolds derived from a known mRNA target (SON), a primary microRNA (pri-miR-30a), and an artificial sequence predicted to lack secondary structure (Fig. 3C; fig. S3, A and B). Different scaffolds affect methylation levels, but the wild-type enzyme consistently modifies GGAC more efficiently than GGAU while the R298P mutant has the reversed substrate preference. Different substrate binding affinities could explain the altered methylation substrate preference, even though SELEX was unable to detect a clear consensus. In electrophoretic mobility shift assays, there are no significant differences between the wild-type and the mutant enzyme in their affinity for RNA substrates containing GGAC or GGAU motifs (fig. S3C). Thus, the observed changes in methylation efficiencies are not due to changed overall affinities for the RNA substrate. Furthermore, the affinity for the methylated RNA is also similar (fig. S3D), suggesting that product release is not the step affected by R298P. Together, quantitative enzymatic assays and binding assays show that METTL14^R298P^ changes the substrate sequence preference for methylation, without affecting binding affinities.

**Fig. 3.**
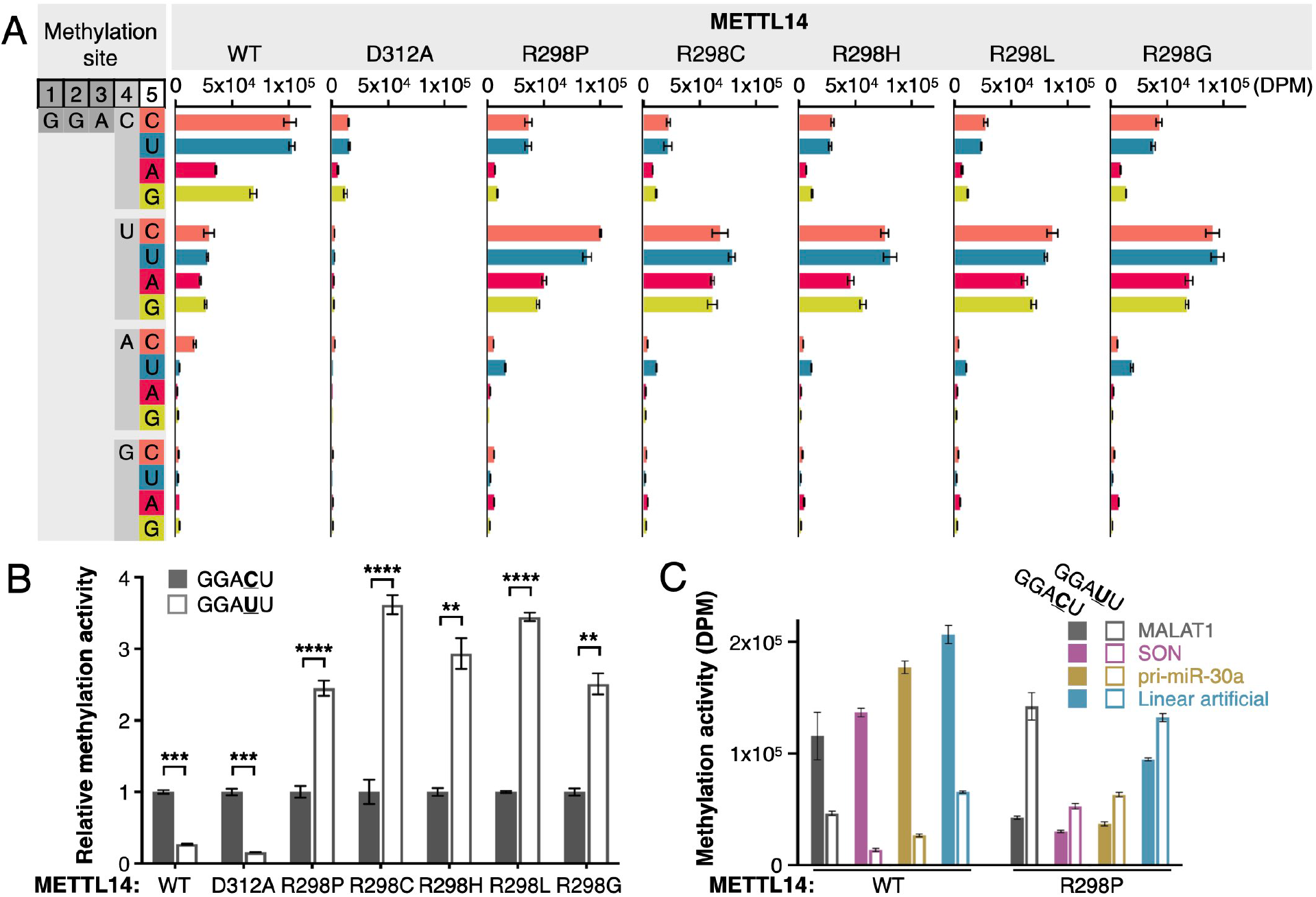
METTL14^R298P^ methylates GGAU with higher efficacy. **(A)** Quantitation of in vitro methylation activity of recombinant METTL3-METTL14 complexes with the indicated METTL14 allele, on a series of RNA fragments derived from MALAT1 (with single nucleotide changes at 4^th^ or 5^th^ positions). Methylation activity is represented with the amount of ^3^H-methyl incorporation measured as disintegration per minute (DPM). Data represented as mean ± s.d. from three replicate reactions. (B) Methylation activity of the GGAUU substrate normalized to that of GGACU for each protein mutant. Data represented as mean ± s.d. from three replicate reactions. Unpaired two-tailed t-test results are indicated with asterisks; **, *P* < 0.01; ***, *P* < 0.001; *”*, *P* < 0.0001. (C) In vitro methylation of RNA substrates containing GGACU or GGAUU in different sequence and structural contexts. Data are represented as mean ± s.d. from three replicate reactions.

### METTL14^R298P^ promotes m^6^A modification at distinct sites in the transcriptome

We postulated that the altered substrate specificity of METTL3-METTL14^R298P^ would change m^6^A modification profiles in mRNAs. To detect the m^6^A sites in polyadenylated RNAs, we adopted the m^6^A-seq workflow (10, 11). While there are other methods that can locate the methylation marks at a higher resolution than m^6^A-seq (e.g. miCLIP, DART-seq, m^6^A-REF-seq) (19–21), most of them rely on the presumed sequence near the modification site—Cyt immediately following the methylated Ade. However, such methods would arbitrarily prevent the detection of modified adenines in different sequence contexts, especially the GGAU motif we found from IVM-seq that is the most preferred substrate for the R298P mutant. Thus, to avoid any bias about the sequence composition of the m^6^A sites, we only relied on immunoprecipitation with antibodies (m^6^A-seq).

From each cell line expressing a unique METTL14 allele, we identified ~5000–7000 total m^6^A-seq peaks that are reproduced in all three biological replicates (Fig. 4A). Each sample usually yields >10,000 peaks per replicate using our stringent peak-calling criteria. The inherent incomplete coverage of the m^6^A-seq method results in a lower number of peaks reproduced in all replicates, but the peak overlap is generally in the expected range (22). Among the tested samples, METTL14^R298P^ peaks seem to overlap less with the peaks found in the other samples (fig. S4A). To dissect the difference among the METTL14 alleles, we identified peaks unique to each cell line by selecting the peaks that do not overlap with the peaks in the other three cell lines in a 4-way comparison (Fig. 4A). The distribution of total and unique peaks is similar within mRNAs and genome-wide (fig. S4, B and C). Consensus motif analysis of the m^6^A peaks unique to each cell line shows that R298P cells have a preference for Ura after the methylated Ade, while the other samples retain the dominant Cyt signal at the same position (Fig. 4B). The preference for Ura in R298P cells is less prominent when all m^6^A sites are analyzed for total m^6^A sites, which may be due to the background from endogenous wild-type METTL14 (fig. S4D).

**Fig. 4.**
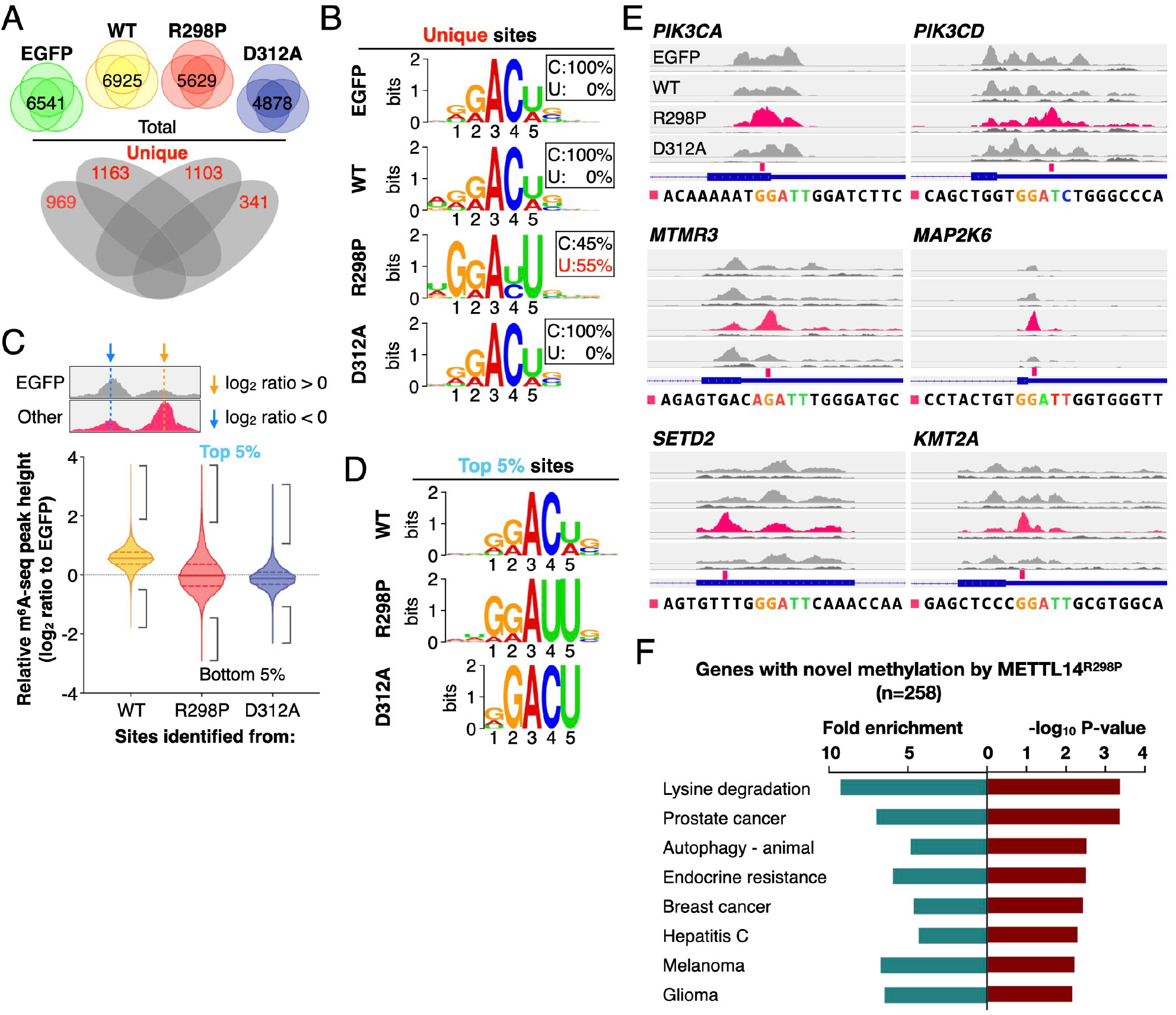
METTL14^R298P^ promotes m^6^A modification of distinct RNA sites in the transcriptome. **(A)** Four stable HepG2 cell lines expressing EGFP control or METTL14 (WT, R298P or D312A) were used for m^6^A-seq analysis. The number of peaks reproduced in all 3 replicates of each sample is shown as intersection on top (total), and the unique peaks after a 4-way comparison are shown below (unique). (B) Motif analysis of the m^6^A sites unique to each cell line (marked with red numbers in (A)). The probability percentages of Cyt and Ura at the 4^th^ position are shown in a box. MEME P value from top to bottom: 2.2e^-119^, 8.7e^-75^, 5.6e^-103^, and 8.8e^-21^. **(C)** Quantification of m^6^A-seq peak height from cell lines expressing METTL14 (WT, R298P or D312A) relative to the EGFP control. Schematic of summit signal comparison workflow is shown on top. Data are shown as median and quartiles with dashed lines (WT: n=33549; R298P: n=28892; D312A: n=267O3). The brackets mark top or bottom 5 percentiles. (D) Motif analysis of the top 5% methylation sites (marked cyan in (C)). MEME Pvalues from top to bottom: 7.4e^-81^, 7.7e^-349^, and 1.1e^-21^. **(E)** Browser tracks of example transcripts with a peak unique to METTL14^R298P^ (pink). The height of each track is normalized by the total count of mappedreads per sample, and the called R298P-unique peaks are indicated with pink tick marks underneath. Immunoprecipitate (top) and input (bottom) track pairs shown for each sample. The sequence around each pink tick mark is shown at the bottom, with the consensus motif highlighted with color. (F) Gene ontology analysis (DAVID) of the genes (n=258) that contain at least one m^6^A peak unique to R298P cells (A) and one tall m^6^A peak (top-5%, relative to EGFP) (C).

To investigate the altered substrate specificity of METTL14^R298P^ further, we used another method in parallel to identify the novel m^6^A peaks that increase in intensity upon expressing the GOF mutant. We compared the height of each m^6^A peak to the signal in the EGFP control (each normalized to sample total read counts) at the same genomic location (Fig. 4C). We then used the sequences with different relative peak heights (top or bottom 5%) for motif analysis (Fig. 4D). For the R298P sample, the tallest peaks relative to the EGFP control (top 5%) showed a clear preference for Ura at the 4th position, but the shorter peaks compared to the EGFP control (bottom 5%) maintain a strong preference for Cyt at the 4th position (fig. S4E). No such difference in sequence preference was observed for the other cell lines. Manual inspection of the m^6^A-seq gene-browser tracks of the “top 5%” peaks provided examples of the peaks that are taller in R298P cells, and we can readily identify the signature sequence preference of the GOF mutant, GGAU (Fig. 4E and fig. S4F). Therefore, the mutant methyltransferase METTL3-METTL14^R298P^ exhibits a novel RNA sequence preference—GGAU rather than GGAC—in the transcriptome. We also performed GO analysis on the most stringent set of R298P-dependent novel m^6^A peaks— the peaks that are taller in R298P (top-5%) and lack overlap with m^6^A peaks from any of the other samples (R298P-unique). The most robust sites targeted by METTL14^R298P^ happen to be involved in histone modification and cancer pathways (Fig. 4, E and F; fig. S4F). How the novel methylation events caused by the GOF mutant methyltransferase lead to altered gene expression important for cancer, including changes in WNT signaling and cell motility needs further investigation and may change with different cellular contexts.

Having confirmed that METTL3-METTL14^R298P^ preferentially methylates a distinct target RNA sequence in vitro and in the transcriptome, we investigated how the changed sequence context may affect m^6^A detection for downstream events. Most m^6^A marks rely on being recognized by a reader protein for effector function via the YTH domain (1, 2). We tested whether the novel methylated sequence, GGm^6^AU, would interfere with binding the reader proteins. Using purified recombinant YTH domains of known readers—YTHDC1, YTHDC2, YTHDF1, YTHDF2, and YTHDF3—we tested how the sequence context affects the affinity of the YTH domains to m^6^A-modified RNA. All the YTH domains bind the methylated RNA more tightly compared to the unmethylated RNA in gel-shift assays, and the affinity for the m^6^A-modified RNA can vary (fig. S5, A to E). Changing the sequence context from GGAC to GGAU did not interfere with the ability of the reader protein to detect the m^6^A modification. Thus, the new m^6^A modifications created by METTL14^R298P^ at GGAU sites that are not typically methylated are likely to recruit m^6^A readers similarly to the canonical m^6^A sites containing GGAC (23).

### Structural basis for altered RNA specificity of METTL14^R298P^

To build a molecular model to understand the novel RNA target specificity of METTL14^R298X^, we determined crystal structures of the methyltransferase domain complexes of METTL3-METTL14, for all three patient-derived mutations of R298 in METTL14 (Table S1). The methyltransferase domains are known to heterodimerize to form an elongated complex where R298 of METTL14 is located > 20 Å away from the bound S-adenosyl methionine (SAM) in the active site of METTL3 (Fig. 5A). In all three mutant structures, the backbones of both proteins maintain a similar conformation as the wild type, but important side-chain rearrangements occur. Superimposing the wild-type structure on that of METTL3-METTL14^R298P^ shows that R471 and H474 of METTL3 move inward to fill the space that is normally occupied by the side chain of METTL14^WT^ R298 (Fig. 5B). For structures containing R298H or R298C mutations in METTL14, we observe electron densities that suggest that both conformations are accessed by the same two residues (Fig. 5C and fig. S6, A to D). Therefore, we conclude that loss of the arginine side chain at 298 allows the protein to access an alternative conformation that preferentially methylates GGAU sequences.

**Fig. 5.**
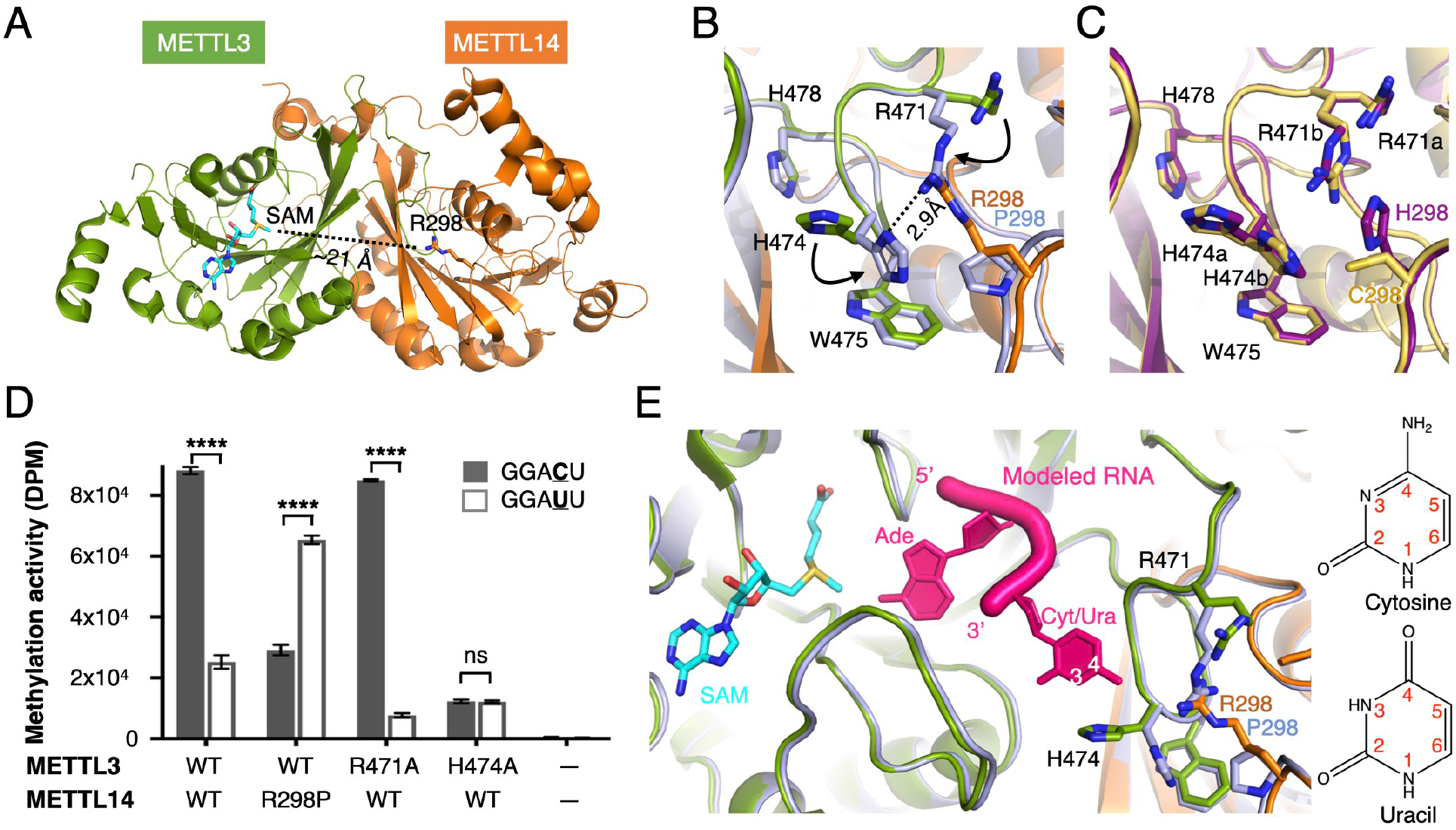
Structural basis for how METTL14^R298P^ gains an alternate substrate specificity. **(A)** Overall structure of wild-type METTL3-METTL14 methyltransferase domains provided for context, and SAM (Cyan) and R298 side chain are shown with stick representation. **(B** and **C)** Close-up views of the METTL3-METTL14 complex crystal structures near the interface. WT (Green and Orange) superimposed on R298P (light blue) (B) and R298C (yellow) superimposed on R298H (purple) (C) in the same orientation. METTL3 residues are shown in black font. For (C), two alternative conformations could be modeled for two residues. (D) In vitro methylation assay of indicated mutant proteins using RNA fragments containing a GGAC or GGAU motif. Data represented as mean ± s.d. from triplicate reactions. Unpaired two-tailed t-test results are indicated with asterisks; ****, *P* < 0.0001; ns, not significant. (E) Model of RNA (hot pink) containing the Ade to be methylated and the 3’ nucleotide bound to superimposed structures of wild-type METTL3 (Green) in complex with METTL14 (Orange) or METTL14^R298P^ (light blue). Positions 3 and 4 (white) of the pyrimidine ring in Cyt or Ura (chemical structures shown on right) may be distinguished via the observed side-chain rearrangements.

Due to the alternative conformations in the mutant structures, we postulated that R471 and H474 in METTL3 are likely to contribute to the changed nucleobase preference at the 4th position. We introduced alanine substitutions and measured the effect on in vitro methylation activity (Fig. 5D). Indeed, both mutations impact the relative preference between GGAC and GGAU substrates, though in opposite directions, indicating that they contribute to recognizing the nucleotide immediately following the methylated Ade. R471 has also been independently found to be mutated in certain cancer patients, raising the possibility that changes in RNA specificity of METTL3-METTL14 may contribute to disease in multiple ways.

Combining our structural and biochemical data, we modeled how RNA substrates bind near the active site of METTL3-METTL14 (Fig. 5E). The methylated adenine can be placed near the SAM binding site by superimposing the RNA-bound structure of another m^6^A writer enzyme, METTL16 (18). Abiding by the RNA geometry constraints, the adjacent nucleotide can be modeled to make direct contacts with H474 and R471 of METTL13 or R298 of METTL14, depending on the sidechain conformations. Cyt and Ura differ at positions 3 and 4 of the pyrimidine ring, and R471 and H474 of METTL3 and R298 of METTL14 together accomplish the normal RNA selectivity profile of METTL3-METTL14. Therefore, while investigating the molecular rationale for the oncogenic phenotype of a GOF mutant, we were also able to gain structural insight into how RNA substrates normally engage with METTL3-METTL14 near the active site.

## DISCUSSION

Our work uncovers how a point mutation in METTL3-METTL14 can transform the RNA methylation substrate specificity and promote oncogenesis. Mutations of METTL14^R298^ induce more methylation of novel sites containing GGAU and less methylation at canonical sites containing GGAC (Fig. 6); using randomized RNA libraries, quantitative enzymatic assays, and transcriptome analyses, we have shown that a single amino acid substitution can alter the RNA modification specificity in vitro and in vivo. The oncogenic phenotype of METTL14^R298P^ is associated with gene expression changes in the WNT/β-catenin signaling pathway and others known to control cell migration and invasion. While the direct targets may depend on cell context, m^6^A modifications of GGAU sites are also likely to be recognized by the YTH family of proteins similarly to the canonical GGAC sites. Combining the biochemical findings and the crystal structures of the METTL14^R298^ mutants, we also propose an atomic model for how substrate RNAs bind METTL3-METTL14 near the catalytic site (Fig. 5E). Thus, we have unveiled the molecular mechanism underlying a cancer mutation as well as the normal recognition of RNA substrates for METTL3-METTL14.

**Fig. 6.**
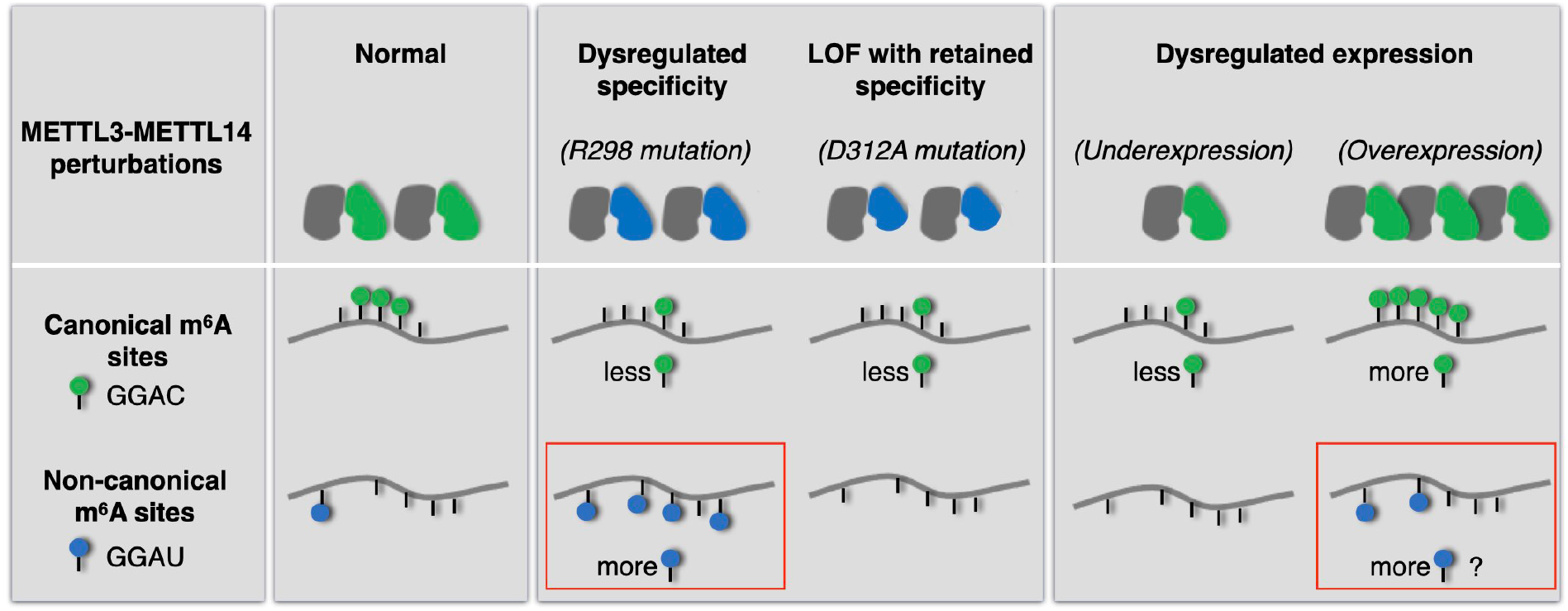
Dysregulated METTL3-METTL14 can cause distinct m^6^A modification states. Our study shows a distinct way to dysregulate METTL3-METTL14 by rewiring its RNA substrate specificity. R298 mutations in METTL14 lead to more methylation of GGAU-containing sites. Increased m^6^A at GGAU sites (red rectangle) is the key difference between R298P and D312A mutations. Thus, more GGAU methylation is linked to the oncogenic phenotype of the R298P mutant. Similar increase of methylated GGAU sites is also likely to occur with overexpression of wild-type METTL3-METTL14, and may contribute to pathogenesis in certain contexts.

Our cancer models show that a change in methylation pattern can promote oncogenesis, without any change in m^6^A levels in mRNAs. Between R298P and D312A (also similar to underexpression case in Fig. 6) mutants, the major difference is the location of m^6^A marks in mRNAs, not the level of m^6^A modification in mRNAs. Therefore, in this context, hypomethylation does not seem to drive oncogenesis; rather, the novel methylation sites created by the R298 mutant are likely responsible for the cancer phenotype. R298 mutations are more frequent than other amino acids within METTL14 in tumor samples, but rarer than many well-known tumor suppressor genes. More commonly, global hypermethylation caused by overexpression of the wild-type METTL3-METTL14 has been linked to multiple cancers. Given that the wild-type methyltransferase has appreciable activity on targets containing GGAU (Fig. 3A and (24, 25)), overexpression may increase methylation of the less preferred sites above a threshold. Thus, in certain cases, overexpression of wild-type METTL3-METTL14 may share a similar pathogenic mechanism as the GOF mutant, METTL3-METTL14^R298P^. Therefore, our study provides a distinct mechanism where noncanonical m^6^A sites containing the GGAU motif should be examined closely in cancers, regardless of how METTL3-METTL14 is dysregulated (by overexpression or mutation).

GOF mutations that change how proteins interact with metabolites, proteins, or DNA have been shown to impact cancer (26–28). Our work shows that a point mutation can rewire an RNA modification enzyme to recognize a distinct nucleotide sequence as the preferred substrate, revealing a powerful GOF mechanism that has not been previously recognized and that may be prevalent in the evolution of cancer cells.

## MATERIALS and METHODS

### Plasmid construction

For co-expression of human METTL3 and METTL14 recombinant protein in *Escherichia coli*, complementary DNA (cDNA) corresponding to full-length METTL3 (a.a. 1-580) and METTL14 (a.a. 1-456) were cloned into pETDuet vector in previous work (4). Point mutations were introduced by site-directed mutagenesis. For lentiviral constructs expressing METTL14^WT^, METTL14^R298P^ and METTL14^D312A^, corresponding sequences were subcloned into the pLJM1-EGFP vector (Addgene plasmid #19319). To express the YTH domain of human YTHDC1 in *E*. coli, the cDNA corresponding to a.a. 345-509 were subcloned into the pET21a vector (Novagen). For the constructs used for mouse hydrodynamic transfection, cDNA corresponding to full-length METTL3 and METTL14 with specific mutations were each subcloned into the pT3-EF1α vector.

### Cell culture

Human liver cancer cell line HepG2 was obtained from ATCC (HB-8065). HepG2 cells were cultured in MEM with EBSS, 2 mM L-glutamine (Cytiva SH30024) supplemented with 1% nonessential amino acids (Gibco 11140-050) and 10% FBS (Corning 35-011-CV) under standard mammalian cell culture conditions (37 °C, 5% CO_2_). The 293 Lenti-X cell line was a gift from D. Nijhawan (UTSW). 293 Lenti-X cells were cultured in DMEM (Sigma D6429) supplemented with 10% FBS under standard mammalian cell culture conditions.

### Virus production and generation of stable overexpression cell lines

To generate lentiviruses for overexpression of EGFP control, METTL14^WT^, METTL14^R298P^ and METTL14^D312A^, each pLJM1 plasmid was co-transfected with psPAX2 (Addgene plasmid #12260) and pMD2.G (Addgene plasmid #12259) using Lipofectamine 3000 (Invitrogen L3000015) into the 293 Lenti-X cells. Viral particles containing media were collected at 72 hours after transfection and filtered by 0.45 μm PVDF syringe filter (Millipore SLHVR33RS). For the generation of stable overexpression of EGFP control, METTL14^WT^, METTL14^R298P^ and METTL14^D312A^ HepG2 cell lines, 2.5 million cells were seeded on each 100 mm plate 24 hours before the viral transduction. The viral transduction was accomplished by incubating cells with a mixture of 10% (v/v) virus-containing media and complete growth media in the presence of Polybrene (10 μg/mL, Sigma TR-1003-G) for 48 hours. Puromycin (3 μg/mL) was added to the growth media for selection. The selection lasted 8 days when fresh puromycin-containing media was provided every 48 hours. Cells were subcultured as needed to maintain the cell confluence between 20% to 80%.

### Quantification of m^6^A by mass spectrometry

Total RNA was extracted from 10 million cells using Trizol (Invitrogen 15596026) according to the manufacturer’s instructions. RNA was digested with DNase I (10 U) at 37 °C for 30 min and extracted by acid phenol-chloroform and precipitated by isopropyl alcohol before dissolving in nuclease-free water. Poly(A) RNA was selected from the total RNA with NEBNext Poly(A) mRNA magnetic isolation module (NEB E7490L), using 100 μL beads slurry per 100 μg total RNA. Poly(A) RNA samples (200 ng) were denatured at 70 °C for 5 min in a 20 μL digestion buffer (50 mM sodium acetate pH 5.5) prior to the incubation with 2 U of nuclease P1 (Sigma N8630) at 42 °C for 2 h. The digested samples were mixed with 3 μL ammonium bicarbonate (1 M), 1 μL MgCl_2_ (25 mM), and 5 U of Antarctic phosphatase (NEB M0289S) then incubated at 37 °C for 2 hours. The samples together with serial dilutions of adenosine (0.05-5 μM, Sigma A9251) and **m^6^A**(1-100 mM, MCE HY-N0086) were analyzed on RapidFire 300 high-throughput SPE system (Agilent Technologies) interfaced with a Sciex 6500 (Sciex). For RapidFire sample preparation, the load/wash solvent (solvent A) was water containing 0.1% trifluoroacetic acid. The elution solvent (solvent B) was acetonitrile/water (8:2, v/v) containing 0.1% trifluoroacetic acid. Samples were aspirated from 384-well plates and loaded onto a SPE cartridge (cartridge type C, C_18_) to remove buffer salts, using solvent A at a flow rate of 1.5 mL/min for 3000 ms. The retained and purified analytes were eluted to the mass spectrometer by washing the cartridge with solvent B at 1.25 mL/min for 5000 ms. The cartridge was re-equilibrated with solvent A for 600 ms at 1.5 mL/min. For spectrometer detection, adenosine and *N*^6^-methyladenosine were measured using a selective reaction monitoring protocol. The parent ions in Q1 and their corresponding daughter ions in Q3 were as follows: (i) Q1 mass = 268.2 amu > Q3 mass = 135.9 amu for adenosine, (ii) Q1 mass = 281.7 amu > Q3 mass = 149 amu for *N*^6^-methyladenosine. The resolution for Q1 and Q3 were set to ‘unit.’ A dwell time of 150 ms was used for each transition. For adenosine detection the declustering potential was set to 25 V, the entrance potential to 10 V, collision energy to 27 V and collision cell exit potential to 16 V. For *N*^6^-methyladenosine detection the declustering potential was set to 36 V, the entrance potential to 10 V, collision energy to 19 V and collision cell exit potential to 4 V. For both nucleosides the ion spray voltage was set to 5500 V and the source temperature was 700 °C. The gas settings were: curtain gas (30 psi ion); source gas 1 (70 psi); ion source gas 2 (70 psi). The area under the daughter ion peaks (area under the curve, AUC) was quantified using RapidFire integrator software. The absolute concentrations of adenosine and *N*^6^-methyladenosine from each sample were determined by the standard curves generated from each serial dilution of the m^6^A and A, and then the m^6^A/A ratios were calculated.

### Cell invasion, migration, and proliferation assays

For HepG2 stable cell line invasion assay, the inserts (Corning 354480) pre-coated with Matrigel were rehydrated with serum-free MEM with 1% non-essential amino acids for 2 hours in the cell culture incubator. 150,000 cells per insert in serum-free MEM media (0.5 mL) were seeded inside the insert and another 0.5 mL complete growth media was placed on the bottom of the well. Following 72-hour of invasion, cells were fixed and stained in the crystal violet solution (5 mg/mL crystal violet, 20% methanol in water solution, filtered) for 30 min. The inserts were extensively washed in water and air-dried. Eight images on different locations of the same insert were taken under the brightfield using an inverted microscope (20x). Images were analyzed by ImageJ (29), and the percentage of areas above the automatically determined threshold was quantified for each image. The average of a total of 8 images was calculated for each sample, and the normalization to the corresponding EGFP control was performed for each batch of experiments. Each batch of experiments at one time was considered a biological replicate. For the cell migration assay, similar procedures were followed except for using a non-coated insert (Corning 353097). The cell invasion and migration images of EGFP samples were only quantified to normalize the data, thus they were not plotted or tested for statistical significance. For the proliferation assay, 5,000 cells per well were seeded on 96-well plates and 3 replica plates were prepared on day 0. The number of viable cells was determined using CellTiter 96 AQueous One Solution Cell Proliferation Assay kit (Promega G3580) and absorbance at 490 nm was measured by a plate reader (CLARIOstar) on day 1, day 3, and day 5. The absorbance values from day 3 and day 5 were normalized to the corresponding absorbance value from day 1 and plotted.

### Mouse Tumorigenesis Assay

All mice were handled in accordance with the guidelines of the Institutional Animal Care and Use Committee at University of Texas Southwestern Medical Center. The Alb-Cre; Lin28a^f/f^; Lin28b^f/f^ and p53^f/f^ mice were on the FVB strain background. Liver cancer inducing oncogenes (pT-CAG/NrasG12V) were co-injected using the Hydrodynamic Transfection (HDT) method with pT3-METTL3^WT^ and pT3-METTL14^WT/R298P/D312A^ along with Sleeping Beauty Transposase (SB100) into 8-week-old mice. Mice body weights were recorded, and livers were harvested six weeks after HDT. Gross livers were weighed and photographed to determine the tumor burden. Mouse livers retrieved from the tail vein injection experiment were fixed in 10% neutral buffered formalin, then dehydrated, cleared, and infiltrated with paraffin according to the standard protocol. 5 μm-paraffin sections were prepared from the processed organs and stained with hematoxylin and eosin (H&E). Images were captured by a bright-field microscope (10 x objective) and independently reviewed by a certified pathologist (E.B) affiliated with the UT Southwestern Histopathology Core.

### m^6^A-seq and data analysis

The procedure was adapted from the original m^6^A-seq or MeRIP protocols (10, 11). Three biological replicates of 40 million (2× 150 mm plates) HepG2 cells of stable overexpression of EGFP, METTL14^WT^, METTL14^R298P^ and METTL14^D312A^ were lysed by Trizol and the total RNA was extracted. The total RNA was treated by DNase I (5 U per 100 μg total RNA) and mRNAs were enriched by NEBNext Poly(A) mRNA magnetic isolation module (NEB E7490L). 8-10 μg of mRNA from each sample was fragmented to the size of ~120-nt by Ambion Fragmentation reagent (Thermo Scientific AM8740) in a proportional volume (10 μL reaction per 1 μg mRNA) at 70 °C for 7 min then precipitated in 75% ethanol with 10% volume of sodium acetate (3 M, pH 5.5) and GlycoBlue (1 μL, Invitrogen AM9515). The RNA fragments were dissolved in nuclease-free H_2_O at a concentration of 100 ng/μL, and 200 ng from each sample was spared as input.

5 μg of RNA fragment was denatured and incubated with 15 μg of anti-m^6^A antibody (Abcam ab151230) in immunoprecipitation (IP) buffer (10 mM Tris pH 7.5, 150 mM NaCl, 0.1% NP-40, 5 mM EDTA pH 8.0, 0.2 U/μL SUPERase·In) for 4 hours at 4 °C. The complex was incubated with 100 μL slurry of protein A/G beads blocked by BSA (0.5 mg/mL in IP buffer) at 4 °C overnight. The beads were washed by 0.9 mL, twice of each IP buffer, high-salt buffer (10 mM Tris pH 7.5, 1 M NaCl, 1% NP-40, 0.5% sodium deoxycholate), low-salt buffer (10 mM Tris pH 7.5, 50 mM NaCl, 0.1% NP-40), and FastAP buffer (10 mM Tris pH 8.0, 5 mM MgCl_2_, 100 mM KCl, 0.02% Triton X-100). The IP samples were dephosphorylated on-bead by 5 U of FastAP (Thermo Scientific EF0615) in 100 μL slurry at 37 °C for 30 min in a thermomixer with 15 s pulse-shaking at 1200 r.p.m. every 5 min, then washed by FastAP buffer and T4 RNA ligase buffer (50 mM Tris pH 7.5, 10 mM MgCl_2_, 1 mM DTT). A linker RNA oligonucleotides purchased from IDT was preadenylated at 5’ -end by T4 RNA ligase I (NEB 0437) using a method described previously (30). The pre-adenylated linker RNA (100 pmol) was added to the 3’ -end of IP RNAs on-bead by 500 U of T4 RNA ligase 2, truncated K227Q (NEB M0351) in 50 μL slurry (with a 25% final concentration of PEG8000) at 16 °C for an overnight-incubation in a thermomixer with pulse-shaking, then washed by T4 RNA ligase buffer and high-salt buffer. The input samples were dephosphorylated and ligated to the pre-adenylated linker RNA similarly in tubes, except the reactions were scaled-down 5-time and 2.5-time, respectively. Precipitation of RNA in 80% ethanol was performed after both reactions for the input samples. The IP RNA on-bead was eluted twice by incubation with 100 μg proteinase K in a 100 μL slurry (50 mM Tris pH 8.0, 50 mM NaCl, 1 mM EDTA, 1% SDS, 0.4 U/μL SUPERase·In) at 50 °C for 1 hour in a thermomixer with pulse-shaking then extracted by acid phenol-chloroform and precipitated in 75% ethanol with GlycoBlue and 0.1× volume of NaCl (2 M). Input and IP RNA samples were reversed transcribed by RT primer using SuperScript III and the excessive RT primer was removed by ExoSAP-IT. The RNA template in RT reactions was removed by NaOH (3 μL, 1 M) hydrolysis at 70 °C for 12 min and the reaction was neutralized by 3 μL HCl (1 M). The resulting first-strand was cleaned by MyONE Silane beads (Invitrogen 37002D) before ligated to an ssDNA adaptor fused with 10-random-nucleotide UMI (60 pmol) by T4 RNA ligase 1 (45 U) in a 30 μL reaction with 20% PEG8000, and cleaned again by MyONE Silane beads after ligation. The standard Nextera i5 and i7 barcodes were added to the adapted cDNA by a 13-cycle of PCR amplification. The dsDNAs were resolved on 2% agarose gel, and the fragments within the 175-250 bp range were eluted and pooled for Illumina NextSeq500 single-end 75-bp sequencing with expected sequencing depth of 25 million reads.

The quality of resulting m^6^A-seq datasets were assessed using the FastQC (version 0.11.2) (http://www.bioinformatics.babraham.ac.uk/projects/fastqc) and FastQ Screen (version 0.4.4) (http://www.bioinformatics.babraham.ac.uk/projects/fastq_screen). The low-quality reads and sequencing adapters were removed by Trim Galore (version 0.4.4) (https://www.bioinformatics.babraham.ac.uk/projects/trim_galore). The reads were aligned to the human reference genome (hg38) using STAR (version 2.7.9a) (31), with the parameters set for aligning reads once. The duplicated alignments were eliminated using Picard toolkit (Broad institute). The resulting primary mapped reads were 6.6-22.3 million per sample. Uniquely mapped reads were called peaks for the IP populations against its input by MACS (version 2.1.2) (32) with additional parameters (--nomodel --extsize 66 -p 1.00e-05). To achieve a higher resolution and stringency, the resulting summits (coordinates of single-nucleotide) from peakcalling were defined as the center of the 50-nt peaks, which is the default peak-length in this m^6^A-seq study. The 50-nt peaks overlapped among three replicates (at least 1-nt with at least one other replicate) derived from the same cell line were assembled as intersectional population using HOMER (version 4.9) (33) mergePeaks function. Each peak summit of newly defined reproducible total populations was the average of three original summit coordinates, which participated in overlap. The peak region of the reproducible total population was expanded to 50-nt around its summit. The peak population unique to each cell line (non-overlapping with any peak from the other three cell lines) was obtained by another round of HOMER mergePeaks analysis. The resulting peak population numbers were summarized (Fig. 4A). The percentage overlap of total m^6^A-seq peaks between any two replicates were calculated during the mergePeaks analysis, and depicted in a series of heatmaps (fig. S4A) to illustrate the sample pairwise-comparison of the peak-overlapping situation. Metagene plots (fig. S4B) depicting the distribution of m^6^A sites across the length of mRNA transcripts were generated by MetaPlotR (34). For the analysis of site distribution on genomic regions (fig. S4C), the location of peaks was annotated by the annotatePeaks module in HOMER. For the discovery of methylation site sequence consensus of the reproducible total population from each cell line (fig. S4D), the coordinates of 50-nt peaks were analyzed by the findMotifsGenome module in HOMER for a *de novo* search. For the discovery of methylation site sequence consensus of the unique population from each cell line (Fig. 4B), the strand information of the summits was retrieved from the source of GTF files on Genecode (35). The actual 50-nt sequences of the peaks were extracted using BEDtools (version 2.17.0) (36) getfasta function with appropriate strand orientation and then analyzed by MEME (version 5.1.1) using the Classic mode. The output position-specific probability matrices from HOMER and MEME analysis were reconstituted to logographs by the Seq2Logo (version 2.0).

For the m^6^A-seq quantitative analysis, the full-union of summits of MACS peak-calling from 3 replicates were designated as the methylation sites index for each cell line. The read-count aligned on the methylation site index was converted from the STAR alignment by the bedcov function in Samtools (version 1.10). To compare the methylation levels of R298P vs. EGFP on the R298P methylation sites, the read-counts of all 3 replicates from EGFP and R298P were determined, converted to CPM, and averaged. The ratio and log_2_ fold-change were calculated and used in the plot (Fig. 4C). The same process was done for the pairwise comparisons of WT vs. EGFP and D312A vs. EGFP. To describe the methylation consensus sequence of the sites with the highest or the lowest relative m^6^A enrichment, 50 nucleotides surrounding the summits were extracted and analyzed by MEME.

For the demonstration of representative transcripts harboring the GGAUU sites (Fig. 4E and fig. S4F), the aligned sequencing reads files derived from input and IP were visualized by Integrative genomics viewer (version 2.8.0) (37) as tracks with a height normalized to count of mapped-reads. m^6^A site enrichment (top track) and input (bottom track) were presented in pairs for each m^6^A-seq population. The corresponding GGAUU sites in transcripts were manually located within 50-nt of the peak region.

### Differential RNA expression analysis

Mapped-reads from three biological replicates of input RNA samples of each cell line in m^6^A-seq experiment were counted by featureCounts module of Rsubread package (version 1.4.6) (38). ComBat-seq was used for correcting batch effects between libraries prepared and sequenced at different times (39). Differential RNA expression analysis in pairwise comparisons between cell lines were performed using edgeR (40). The transcripts with zero count were eliminated and the TPM values from three replicates were used to calculate log2 fold change and FDR. Gene ontology analysis was performed using Ingenuity Pathway Analysis tool (Qiagen) and DAVID (41)

### Online 2D LC-MS/MS analysis

Approximately 2×10^6^ cells were lysed by incubating in 120 μL of CelLytic M (Sigma C2978) containing protease inhibitor cocktails on ice for 30 min. The cell lysates were centrifuged at 16,000 ×g for 30 min at 4°C to remove cell debris. The proteins were quantified by using Bradford assay and subsequently subjected to filter-aided sample preparation using trypsin as the digestion enzyme, as described previously (42). In brief, 80 μg of total proteins were denatured in a buffer containing with 8 M urea and 50 mM NH_4_HCO_3_ in a 30 kDa molecular weight cut-off polyethersulfone membrane centrifugal filter unit (VWR 82031-354). Cysteines in the denatured proteins were reduced and alkylated by incubating with 20 mM DTT at 37°C for 1 hour and 55 mM iodoacetamide in the dark at room temperature for 20 min. The samples were washed three times with 50 mM NH_4_HCO_3_ to remove urea and excess reagents before digested with 2 μg of trypsin at 37°C for 18 hours, and the resulting tryptic peptides were desalted with C_18_ tips (Thermo Scientific 87784). For tandem mass tags (TMT) isobaric labeling, 10% of the resulting peptides were reconstituted in 8.5 μL of 50 mM HEPES pH 8.5. Approximately 4% of the TMTsixplex reagents (Thermo Scientific 90068) were added and the resulting mixtures were vortexed at room temperature for 2 hours. The reactions were quenched by 0.5% final concentration of hydroxylamine and the mixture was further vortexed at room temperature for an additional 15 min. To each sample was added 8.92 μL solution of water/ACN/formic acid (8/1/1, v/v), and the mixture was desalted with C_18_ tips.

LC-MS/MS analysis of TMT-labeled peptides was conducted on an Orbitrap Fusion Lumos tribrid mass spectrometer (Thermo Scientific) coupled with an Easy-nLC 1200 UPLC system (Thermo Scientific). The mass spectrometer was equipped with a high-field asymmetric-waveform ion mobility spectrometry (FAIMS), where the compensation voltages (CV) were set at −40, −60, and −80 V. The carrier gas flow was set at 4.2 L/min. The cycle time was 3 sec with each CV being scanned for 1 sec. The online 2D LC was conducted as described previously (43). In brief, desalted peptides were reconstituted in 10 mL of 5 mM ammonium formate/ 5% acetonitrile and loaded onto a 3-cm capillary strong cation exchange (SCX) column (150 μm i.d.) packed in-house with Luna SCX resin (5 μm in particle size and 100 Å in pore size, Phenomenex) with buffer A (0.1% formic acid in water) at a flow rate of 3 mL/min. The peptides were sequentially eluted by a concentration-series of 5 mL plugs of ammonium formate/acetonitrile solutions. The concentration-series are: (mM of ammonium formate/ % of acetonitrile) #1, 5/5; #2, 70/5; #3, 100/5; #4, 150/5, #5, 200/5; #6, 500/5; #7, 200/10; #8, 200/15; #9, 200/20; #10, 250/5; #11, 300/5; #12, 350/5; #13, 200/25; #14, 1000/5. The peptides eluted from the SCX column were loaded onto a 3-cm capillary C_18_ column (150 μm i.d.) packed in-house with C_18_ resin (5 μm in particle size and 120 Å in pore size, Dr. Maisch GmbH HPLC) with buffer A (0.1% formic acid in water) at a flow rate of 3 mL/min. The peptides were eluted from the C_18_ trapping column and separated on a ~25-cm analytical column (5 μm in particle size and 120 Å in pore size, Dr. Maisch GmbH HPLC) packed in-house with C_18_ resin (3 μm in particle size and 120 Å in pore size, Dr. Maisch GmbH HPLC) at a flow rate of 300 nL/min using a linear gradient of 5-37% buffer B (80% acetonitrile/ 0.1% formic acid in water) over 210 min. Eluted peptides were ionized with a Flex nanoelectrospray ion source (Thermo Scientific). The capillary inlet temperature was set at 305°C and the spray voltage was set to 2 kV. Full-scan in the range of *m/z* 400-1500 were acquired at a resolution of 50 k. Maximal injection time and the AGC were set as default for full-scan MS. For MS/MS acquisition, precursor ions were isolated at a window of 0.5 Th and subsequently fragmented by higher-energy collisional dissociation (HCD) at a normalized collisional energy of 38. Fragment ions were scanned at a resolution of 50 k.

Raw LC-MS/MS files were converted to mzXML format and processed in MaxQuant (version 2.0.1.0) (44). Methionine oxidation and N-terminal acetylation were set as variable modifications; cysteine carbamidomethylation was specified as a fixed modification. The type of LC-MS run was set to ‘Reporter ion MS2” with “6plexTMT” as isobaric labels. Reporter ion mass tolerance and the mass tolerance for MS/MS were set at 0.01 Th and 20 ppm, respectively. Mass tolerance for full-scan MS was set as 20 and 4.5 ppm for the first and main search, respectively. Two missed cleavages were allowed for trypsin. Peptide spectra were searched against target-decoy UniProt human proteome database (UP000005640) and the proteins were subsequently filtered at 1% false discovery rate. A minimum ratio determined from 2 peptides was used for protein quantification. The potential contamination and the decoys were removed from the output files. The reporter ion intensities of each protein were normalized against the mean within the samples. The peptide-spectrum match (PSM) counts derived from the identical protein were aggregated and used as input for differential expression analysis by Limma (42). Genes filtered by adj. P value < 0.05 were subjected to pairwise IPA analysis where R298P was compared to others.

### In vitro methylation sequencing (IVM-seq)

*A* degenerate DNA oligonucleotide mixture (Table S2) consisting of 20-random-nucleotide (N20) and flanking constant regions was synthesized by MilliporeSigma with manual adjustment to achieve the overall equal ratio of nucleotides. An initial double-stranded (ds)DNA library was produced by a 7-cycle PCR amplification in a 1 mL reaction containing 20 pmol of N20 oligonucleotides as a template to preserve the designed library complexity. The dsDNA was then gel-eluted and transcribed into a randomized RNA library. The RNA library (6.25 μM) was methylated by the target recombinant methyltransferase heterodimer (0.25 μM) in a 20 μL reaction (50 mM Tris pH 7.5, 0.01% Triton X-100, 15 mM NaCl, 1 mM DTT, 1% glycerol, 5 μM S-(5’-Adenosyl)-L-methionine (Sigma A7007), 20 U SUPERase·In (Invitrogen AM2694)) at room temperature for 2 hours. The methylated RNA library was extracted by acid phenol-chloroform and precipitated with isopropyl alcohol before being reconstituted in nuclease-free water.

The purified RNA library was incubated with 7 μg of antibody against m^6^A (Abcam ab151230 or SySy 202003) in 100 μL of low-salt binding buffer containing 50 mM Tris pH 7.5, 150 mM NaCl, 0.1% NP-40, 50 U SUPERase·In, and bound to protein A/G beads (Thermo Scientific 88802). The mock control sample was prepared by the same procedures but incubated with beads in the absence of antibody. The protein A/G beads were washed with 0.9 mL of low-salt binding buffer once and high-salt buffer (50 mM Tris pH 7.5, 500 mM NaCl, 0.1% NP-40) twice, then one more time by the low-salt binding buffer before the elution by 0.1 M glycine pH 2.5. Methylated RNA species were recovered from the elution and reverse transcribed (RT) using the RT primer and SuperScript III (Invitrogen 18080044). Excessive RT primer was digested with ExoSAP-IT (Applied Biosystems 78250). The resulting cDNA was adapted for barcoding by 10-cycle of PCR amplification and the dsDNA was purified by PureLink PCR Micro kit (Invitrogen K310010). The standard Nextera i5 and i7 barcodes were added to the dsDNA by another 5-cycle of PCR amplification before pooling for Illumina NextSeq500 single-end 75-bp sequencing. The sequencing was performed at Next Generation Sequencing Core, Eugene McDermott Center, University of Texas Southwestern Medical Center. The 20-nt and 21-nt constant flanking sequences from the N20 oligonucleotides were used to trim the reads by using the Trim Ends module in Geneious (version 2021.2.2). The reads with the length of 20-nt after trimming were selected for motif analysis by MEME (45) using the Differential Enrichment mode with the mock control set as background. The output position-specific probability matrices were reconstituted to logograph by the Seq2Logo (version 2.0) (46).

### SELEX

The experiment was performed similarly as described previously (16). An aliquot (0.2 nmol) of randomized RNA library produced for IVM-seq experiment was incubated with equal molarity of 6x His-tag fused target protein complex immobilized on 7 μL Ni-NTA magnetic agarose beads (Thermo Scientific 78605) in the SELEX binding buffer (10 mM Tris pH 8.0, 150 mM NaCl, 10 mM MgCl_2_, 0.01% NP-40, 1% glycerol, 1 mM ß-mercaptoethanol, 10 U SUPERase·In) at 22 °C for 30 min on a thermomixer with 15 s pulse-shaking at 1200 r.p.m. every 5 min. After 3x wash by 180 μL SELEX binding buffer, the RNA species with affinity was recovered from beads by heating at 65 °C for 5 min in 1 mM Tris pH 7.5 buffer containing 20 pmol of RT primer (same as used in IVM-seq). The elution of RNA was assembled in an RT reaction similarly as in IVM-seq. The subsequent cDNA was amplified by dsDNA amplification primers 100 μL PCR reaction to generate the dsDNA template for the next cycle of SELEX. Five SELEX cycles were performed before generating the library for high-throughput sequencing. The SELEX samples were adapted and sequenced together with IVM-seq samples. The sequence data were processed, and motifs analysis was performed identically as for IVM-seq.

### *In vitro* methylation assay

Purification of recombinant full-length METTL3-METTL14 protein complexes was carried out as described for the wild-type construct (4). The RNA oligonucleotides used for methylation activity screen (fig. S1D; Fig. 3, A and B; Fig. 5D) were derived from the context of *MALAT1* methylation site 2577 with single-nucleotide substitutions of the 4^th^ or 5^th^ position within the GGACU site (Table S2). The RNA oligonucleotides and substitutions derived from SON, pri-miR-30a, and the linear artificial RNA (Fig. 3C) are also shown in Table S2. All RNA oligonucleotides were purchased from MilliporeSigma or IDT. The *in vitro* methylation assay was carried out as previously described (4). All reactions were performed in triplicates and the statistical tests used are described in the figure legends.

### Crystallization and Structure Determination

The mutant methyltransferase domain complexes were expressed by introducing the mutation using QuickChange site-directed mutagenesis into the wild-type methyltransferase domain co-expression construct described in (4). Protein purification and crystallization were the same as for the wild type. Crystals were grown by using the hanging-drop vapor-diffusion method by mixing 1 μL protein (15 mg/mL) with 1 μL reservoir solution containing 0.1 M Tris (pH 8.0) and 20% PEG-3350 and incubating at 18 °C. The datasets were collected at APS-19-ID at wavelength 0.9794 Å. Data were indexed, integrated, and scaled by the program HKL3000 (47). Initial phases were obtained by molecular replacement using the wild-type MTD3/MTD14 complex structure (PDB: 5K7M) as a search model. The model was further built manually with COOT (48) and iteratively refined using phenix.refine (49). The PROCHECK program was used to check the quality of the final model, which shows good stereochemistry according to the Ramachandran plot (50). All structure figures were generated by using the PyMOL (Schrodinger, LLC). The software used in this project was curated by SBGrid (51).

### Electrophoretic mobility shift assay (EMSA)

To determine the affinity of METTL3-METTL14 complexes for the GGACU and GGAUU consensus sites (fig. S3, C and D), RNA oligonucleotides derived from the *MALAT1* sequence were used. RNA oligonucleotides were labeled at 5’ ends with ^32^P using T4 polynucleotide kinase and annealed by incubating at 70 °C for 10 min and then snap-cooling on ice. METTL3-METTL14 (0.008-2.05 μM) complexes were incubated with each RNA (1 nM) in a binding buffer (25 mM Tris pH 8.0, 150 mM NaCl, 1 mM DTT, 5 mM KCl, 5 mM EDTA pH 8.0, 1% glycerol, 0.01% Tween-20, 2 ng/μL yeast tRNA) before being separated on native polyacrylamide gels. The YTH domains of YTHDC1, YTHDC2, YTHDF1, YTHDF2, and YTHDF3 were overexpressed in Rosetta (DE3) pLysS cells (Novagen) and purified using its affinity for Ni-NTA agarose (Qiagen). The eluate was further purified by ion-exchange and size-exclusion chromatography steps. RNA oligonucleotides incorporating a single **m^6^A** at the indicated locations were purchased from IDT (Table S2). EMSA was performed similarly as METTL3-METTL14, except for the binding buffer (20 mM Tris pH 8.0, 150 mM NaCl, 10 mM DTT, 50 μM ZnCl_2_, 3.33 ng/μL yeast tRNA, 5% glycerol). Gels were dried and visualized using a phosphor screen and PharosFX Plus imager (BioRad).

### Western blotting

One million cells were washed in Dulbecco’s phosphate-buffered saline (DPBS) and lysed in 100 μL of RIPA buffer (50 mM Tris pH 8.0, 150 mM NaCl, 5 mM EDTA pH 8.0, 1% NP-40, 0.5% sodium deoxycholate, 0.1% SDS). The total protein concentration from the lysate was measured by Rapid Gold BCA protein assay kit (Thermo Scientific A53225) to normalize the amount of total protein in each sample. Blotted PVDF membranes were blocked with skim milk (5%) in TBS-T (20 mM Tris pH 7.6, 150 mM NaCl, 0.05% Tween 20) for 1 hour and incubated with primary antibodies (anti-METTL3, Bethyl A301-567A, 1:1,000 (v/v); anti-METTL14, Thermo Scientific PA5-58204, 1:1,000 (v/v); HRP conjugated anti-β-Actin, Sigma A3854, 1:10,000 (v/v)) overnight at 4 °C with agitation. Each membrane was then incubated with the appropriate HRP-conjugated secondary antibodies in TBS-T and visualized by using the Enhanced chemiluminescence substrate (BioRad 170-5060) and ChemiDoc (BioRad).

### Quantitative PCR

Three biological replicates of HepG2 cells of stable overexpression of EGFP, METTL14^WT^, METTL14^R298P^ and METTL14^D312A^ were lysed by Trizol. The total RNA was extracted and treated by DNase I. cDNA was synthesized from 5 μg of total RNA using oligo dT18 primer (5 μM) and SuperScript III (Thermo Scientific 18080044). Quantitative PCR was performed using the gene specific primer sets (Table S2) on BioRad CFX-384.

### Statistics and reproducibility statement

The western blot of HepG2 stable cell lines were performed twice with similar results. The determination of the m^6^A to adenosine ratio of RNAs from HepG2 stable cell lines was carried out for three biological replicates. Significant difference among means of m^6^A to adenosine ratio in total RNA (fig. S1E) is not found (*P* = 0.3133, ordinary one-way ANOVA). Significant difference among means of m^6^A to adenosine ratio in poly(A) RNA (fig. S1F) is found (*P* < 0.0001, ordinary one-way ANOVA). The adjusted *P* values from post hoc Tukey’s multiple comparison test are: *P* = 0.1667 (EGFP vs. WT, not significant), *P* = 0.0011 (WT vs. R298P), *P* = 0.0076 (WT vs. D312A), *P* = 0.4019 (R298P vs. D312A, not significant). In cell migration and invasion experiments, four or five independent assays were performed at different times. The microscopic images within each figure panel were from the same representative assay. The data plotted for the quantification (Fig. 1, A and B) were statistically tested by an ordinary one-way ANOVA test and the difference among means was found (*P* = 0.0008 (Fig. 1A); *P* < 0.0001 (Fig. 1B)). The adjusted *P* values from post hoc Tukey’s multiple comparison test in Fig. 1A are: *P* = 0.0006 (WT vs. R298P), *P* = 0.0083 (R298P vs. D312A), *P* = 0.2004 (WT vs. D312A, not significant); in Fig. 1B, *P* < 0.0001 (WT vs. R298P), *P* = 0.0001 (R298P vs. D312A), *P* = 0.7936 (WT vs. D312A, not significant). The cell proliferation experiment was performed in three biological replicates. In the mice tail injection experiment, the injection of METTL14^WT^ and METTL14^R298P^ constructs were tested in three cohorts (WT, n=14; R298P, n=10 with two deceased on 35- and 38-day post-injection), whereas the METTL14^D312A^ construct was tested in two cohorts (n=7) at different times. Data of liver to body ratio were statistically tested by ordinary one-way ANOVA test (*P* = 0.0003). The adjusted *P* values from post hoc Tukey’s multiple comparison test are: *P* = 0.0005 (WT vs. R298P); *P* = 0.0025 (R298P vs. D312A); *P* = 0.9927 (WT vs. D312A, not significant). The liver tumor tissue images presented in Fig. 1E are representatives of two different cohorts. The photomicrographs (Fig. 1F) of histological analysis are the representatives from livers of two mice per treatment. The m^6^A-seq experiment was performed with three biological replicates at two different times. In the motif analysis of the m^6^A peaks identified through m^6^A-seq, the *P* values of the consensus for total sites (fig. S4D) are: 1e^-609^ (EGFP), 1e^-604^ (WT), 1e^-491^ (R298P), 1e^-412^ (D312A); for unique sites (Fig. 4B): 2.2e^-119^ (EGFP), 8.7e^-75^ (WT), 5.6e^-103^ (R298P), 8.8e^-21^ (D312A); for the top 5% m^6^A enriched sites (Fig. 4D): 7.4e^-81^ (WT), 7.7e^-349^ (R298P), 1.1e^-21^ (D312A); for the bottom 5% m^6^A enriched sites (fig. S4E): 4.9e^-80^ (WT), 4.0e^-174^ (R298P), 8.2e^-133^ (D312A). The RNA differential expression analysis was performed using three independent replicates derived from m^6^A-seq input samples. Quantitative PCR was performed using three biological replicates. Each Cp value used to calculate the ΔCp is the average of three technical replicates, and the mean ΔΔCp from three biological replicates is shown. Significant differences among means of normalized ΔΔCp is found from multiple unpaired t tests (Holm-Šídák method). The adjusted *P* values are: (from left to right) 0.000067, 0.003577, 0.022717, 0.014916, 0.010704, 0.014916, 0.022717, 0.008168, 0.016658, 0.005696, 0.000018, 0.003577 in fig. S2C; <0.000001, 0.000005, 0.003047, 0.009442, 0.005452, 0.009442, 0.001157, 0.005290 in fig. S2D. The high-throughput proteomics mass spectrometry was performed using 3 biological replicates in different times. The IVM-seq experiment for the identification of methylation site consensus sequence of METTL14^WT^ and METTL14^R298P^ was repeated by using different anti-m^6^A antibodies. The *P* value of the consensus logographs generated by MEME analysis are: 5.2e^-412^ (WT, SySy), 1.5e^-32269^ (WT, Abcam), 4.0e^-355^ (R298P, SySy), 9.1e^-45993^ (R298P, Abcam). The IVM-seq for METTL14^R298C^, METTL14^R298H^, and METTL14^D312A^ was performed by using only the SySy anti-m^6^A antibody. The *P* value of the consensus logographs are 6.5e^-1771^ (R298C), 6.9e^-1094^ (R298H), 4.6e^-7^ (D312A). The *in vitro* methylation assays reported in this study were all carried out in triplicate reactions at the same time. The same assay has been performed by different people more than three times with similar results. The data from the comparison of methylation activity with GGACU and GGAUU sites have been statistically tested by an unpaired two-tailed t-test. The *P* values in Fig. 3B are (from left to right) *P* = 0.000131, *P* = 0.000867, *P* = 0.000080, *P* = 0.000045, *P* = 0.002601, *P* = 0.000095, *P* = 0.001398. The EMSA experiments were performed more than three times with similar results. In the SELEX experiment performed to determine the METTL3-METTL14 binding preference, the *P* value of the top consensus logographs generated by MEME analysis are: 3.7e^-1045^, 1.5e^-894^, 1.3e^-8826^ (WT); 2.0e^-899^, 1.1e^-5651^, 2.5e^-10385^ (R298P).

## Acknowledgments

We thank the Structural Biology Laboratory at UT Southwestern for help with synchrotron data collection. The use of the SBC 19ID beamline at Advanced Photon Source is supported by the United States Department of Energy contract DE-AC02-06CH11357. We thank Melissa McCoy and Bruce Posner in the High-throughput Screening Core at UT Southwestern for help with mass spectrometry experiments. All deep sequencing data were obtained through the Next Generation Sequencing Core at the McDermott Center for Human Growth and Development at UT Southwestern. We are grateful for the helpful comments on genomic data analysis from Dr. Blerta Xhemalce. pLJM1-EGFP plasmid was a gift from David Sabatini. pMD2.G and psPAX2 were a gift from Didier Trono. Y.N. is a Southwestern Medical Foundation Scholar in Biomedical Research (Endowed Scholar Program at UT Southwestern), a Pew Scholar in the Biomedical Sciences, and a Packard Fellow. This study was supported by CPRIT (RP190259 to YN), NIH NIGMS (R01GM122960 to YN), NIH NCI (R01CA258589 to YN), NIH NIEHS (R35 ES031707 to YW), and Welch Foundation (I-2115-20220331 to YN).

## Author contributions

CZ generated the stable cell lines and prepared sequencing libraries. LT and CZ performed the cell growth experiments, and BE performed pathological analysis of the mouse tumors. MH performed the mouse tumorigenesis with the hydrodynamic tail vein assay, supervised by HZ. PW determined and refined the crystal structures. CZ, KAD, EH, and BBK purified proteins and performed the in vitro methylation assays, and CZ and OK carried out gel-shift assays. AK, XZ, and CX performed bioinformatics analysis. RS performed preliminary bioinformatics analysis. YY performed proteomics analysis, supervised by YW. CZ and YN compiled and analyzed the results and wrote the manuscript with input from all co-authors.

## Data and materials availability

The atomic coordinates and the structure factors have been deposited in the Protein Data Bank under accession numbers 7RX7, 7RX8, and 7RX6, for the crystal structures of the methyltransferase domains of METTL3-METTL14^R298P^, METTL3-METTL14^R298H^, and METTL3-METTL14^R298C^, respectively. The m^6^A-seq data have been deposited to the Gene Expression Omnibus database (GSEXXX) and the proteomics data have been deposited to proteomeXchange (PXD039447). All other data are included in the main text or the supplementary materials. Reagents generated in this study are available from the corresponding author with a completed Materials Transfer Agreement.

---

**Fig. S1.**
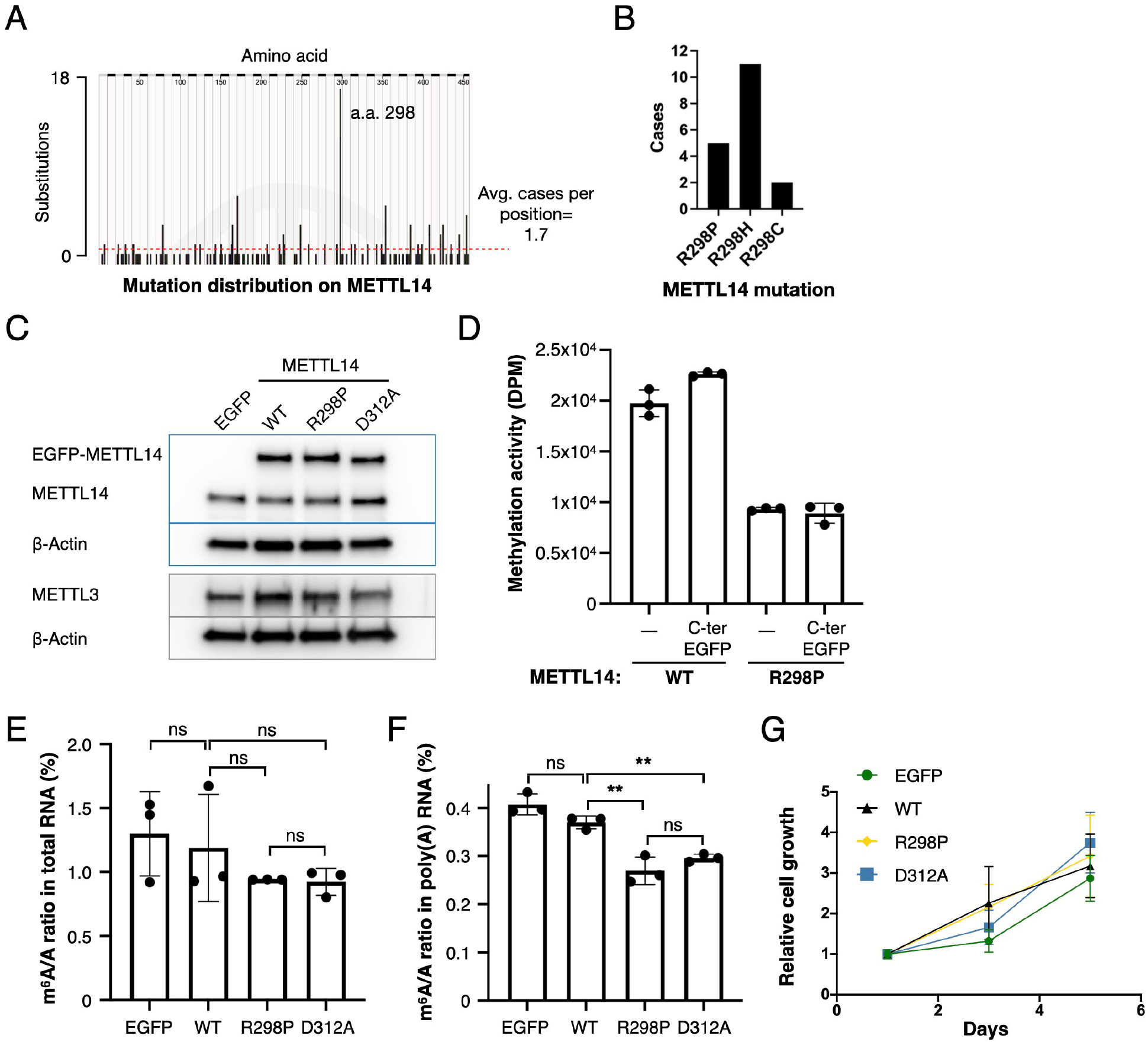
METTL14^R298P^, a mutation found in cancer patients, lowers mRNA m^6^A levels. **(A)** Amino acid mutation frequency of METTL14 in cancer patient genomes. (**B**) Case numbers of various types of cancer carrying R298 mutations of METTL14. (A) and (B) are generated using the data from Catalogue of Somatic Mutations in Cancer (COSMIC, https://cancer.sanger.ac.uk/cosmic). **(C)** Western blot of HepG2 stable cell lines expressing EGFP, EGMETTL14^WT^, METTL14^R298P^, or METTL14^D312A^. The EGFP-fused METTL14 is expressed via viral transduction. (**D**) Quantitation of *in vitro* methylation activity of recombinant METTL3-METTL14 (with indicated METTL14 alleles) on a GGACU-containing RNA fragment derived from *MALAT1.* Data represented as mean ± s.d. from triplicates, with the amount of incorporated ^3^H-methyl measured as the disintegration per minute (DPM). (**E** and **F**) The m^6^A/A molar ratio (%) of total RNA (E) or polyadenylated RNA (F) from HepG2 stable cell lines expressing EGFP, METTL14^WT^, METTL14^R298P^, or METTL14^D312A^determined by LC/MS mass spectrometry. Data are shown as mean ± s.d. from 3 biological replicates. Significant difference among means is found from ordinary one-way ANOVA test in (F) but not in (E). Adjusted P values are indicated: ns, not significant; **, *P* < 0.01 (Tukey’s multiple comparison test). **(G)** Cell proliferation assay using all four HepG2 stable cell lines. The graph is the quantification of viable cells normalized to values measured on day 1. Data are mean ± s.d. from 3 biological replicates.

**Fig. S2.**
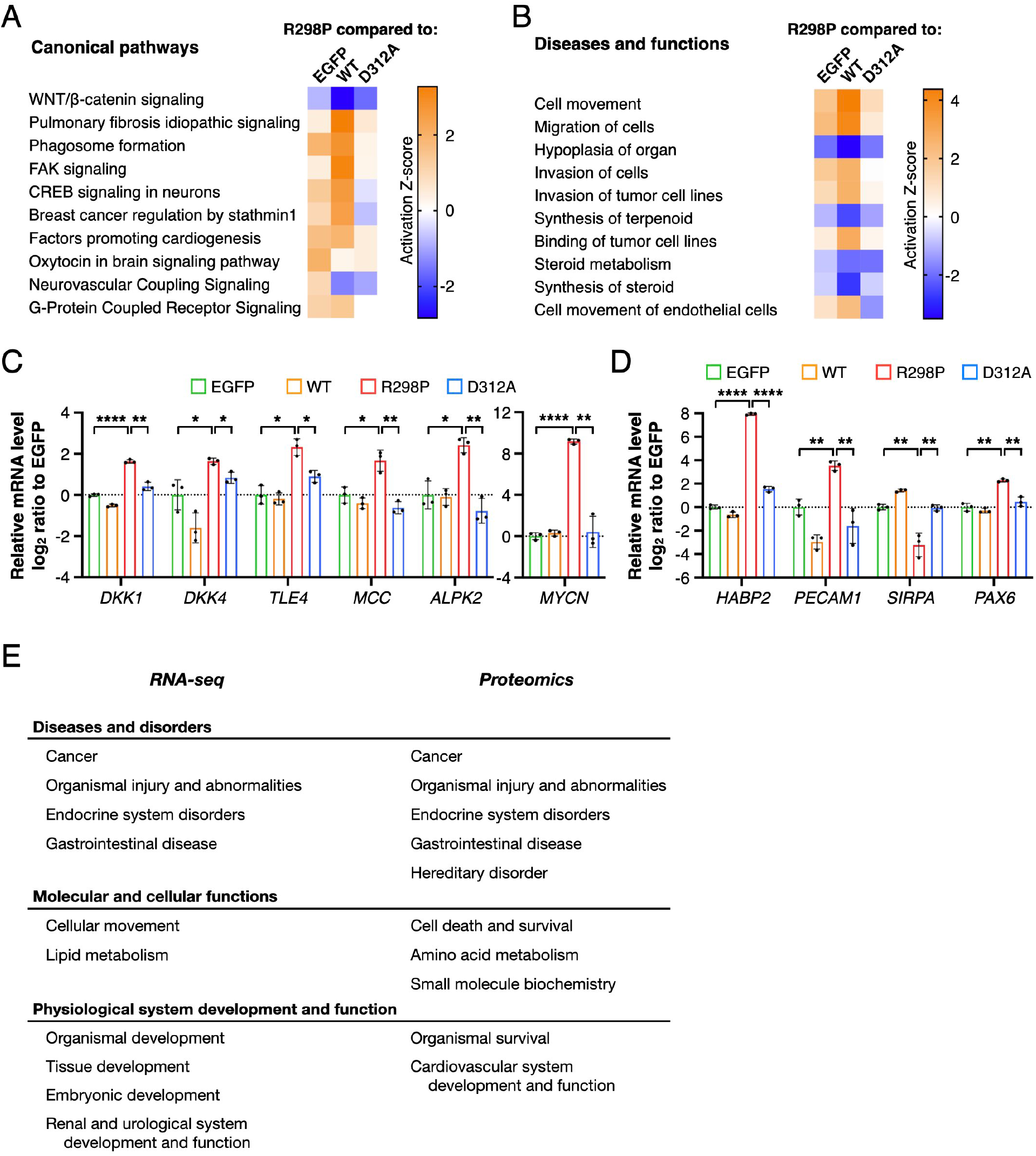
Gene ontology analysis of differentially expressed RNAs and proteins in cells expressing METTL14^R298P^. **(A and B)** RNA-seq differential expression analysis by Ingenuity Pathway Analysis (IPA) Canonical pathways (A) and Diseases and functions (B) modules. Each GO term for each comparison is indicated with a z-score (positive for activation or negative for inhibition in R298P cells). (**C** and **D**) Quantitative RT-PCR of differentially expressed transcripts in METTL14^R298P^ cells. Genes related to the WNT pathway are in (C). Genes involved in cell movement are in (D). Data shown as mean ± s.d. of normalized ΔCp values from 3 biological replicates. Unpaired multiple t-test results are indicated with asterisks; *, *P* < 0.05; **, *P* < 0.01; *P* <0.0001. (**E**) The common among the top-5 GO clusters differentially expressed in R298P (in 3 comparisons of R298P vs EGFP, WT or D312A) using IPA on RNA-seq and proteomics data.

**Fig. S3.**
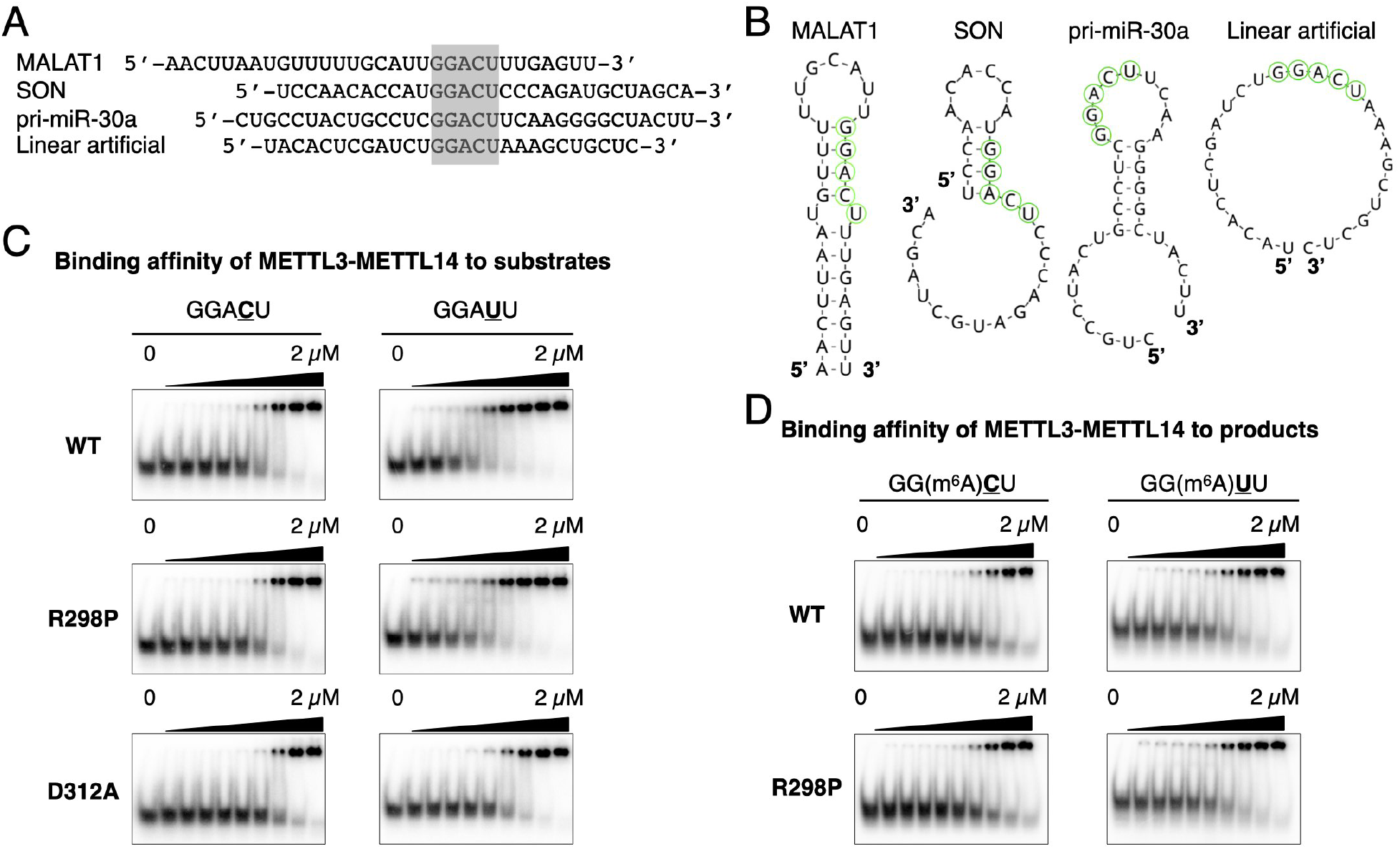
METTL14^R298P^ preferentially methylates GGAU-containing sequences through a mechanism independent of overall RNA affinity. **(A)** The oligonucleotide sequences derived from *MALAT1*, *SON*, pri-miR-3Oa, and a linear artificial sequence containing GGACU (grey). (**B**) Predicted RNA secondary structures {Mathews, 2004 #7250} of the sequences in (A). The methylation consensus sequences are highlighted with green circles. (**C** and **D**) Electrophoretic Mobility Shift Assay of METTL3-METTL14 and RNA oligonucleotides substrates (C) and methylated products (D). Recombinant METTL3-METTL14 concentrations are, from left to right, 0 μM, 0.008 μM, 0.016 μM, 0.032 μM, 0.064 μM, 0.128 μM, 0.256 μM, 0.512 μM, 1.02 μM and 2.05 μM.

**Fig. S4.**
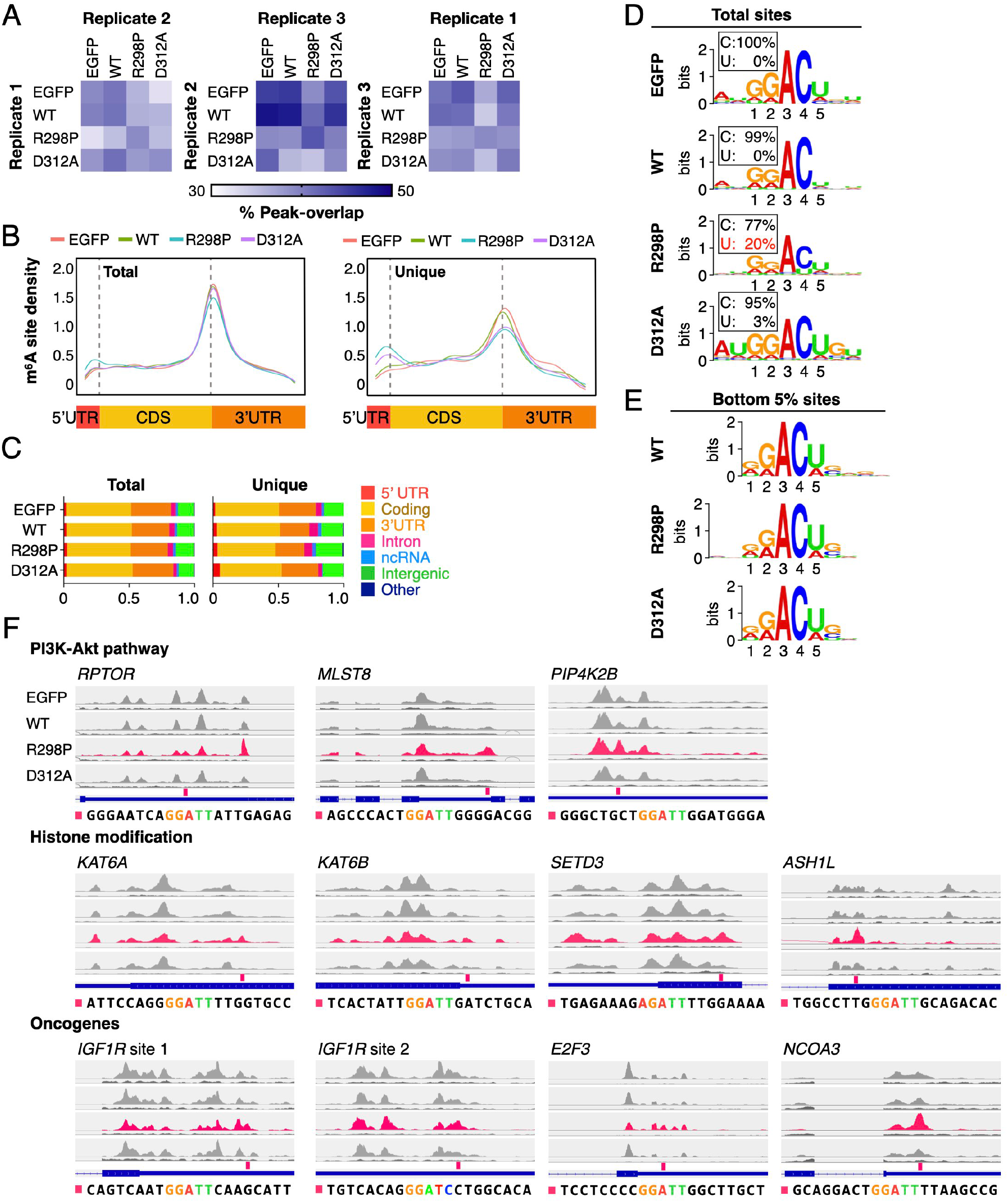
METTL14^R298P^ creates novel m^6^A sites containing GGAUU in the transcriptome. **(A)** Overlap of total m^6^A-seq peaks between each cell line and each replicate. The color gradient represents the % overlap among the called peaks. *, the most peak overlapping of R298P. (**B**) Metagene plots depicting the distribution of total (left) and unique (right) m^6^A-seq peaks across the length of mRNA transcripts. (**C**) The genomic distribution of total and unique m^6^A peaks of each sample reproduced in all 3 biological replicates. **(D)** Motif analysis of the total reproducible m^6^A sites from each cell line. The probability percentages of Cyt and Ura at the 4^th^ position are shown in boxes. The Pvalues of the consensus analyzed by HOMER are (top to bottom): 1e^-609^, 1e^-604^, 1e^-491^, 1e^-412^. **(E)** Motif analysis of the m^6^A sites with the shortest (bottom 5%) peaks relative to EGFP control. The Pvalues of the consensus were analyzed by MEME (top to bottom: 4.9e^-80^, 4.0e^-174^, 8.2e^-133^). (**F**) Browser tracks of example transcripts with a peak unique to METTL14^R298P^(pink), in addition to Fig. 4E.

**Fig. S5.**
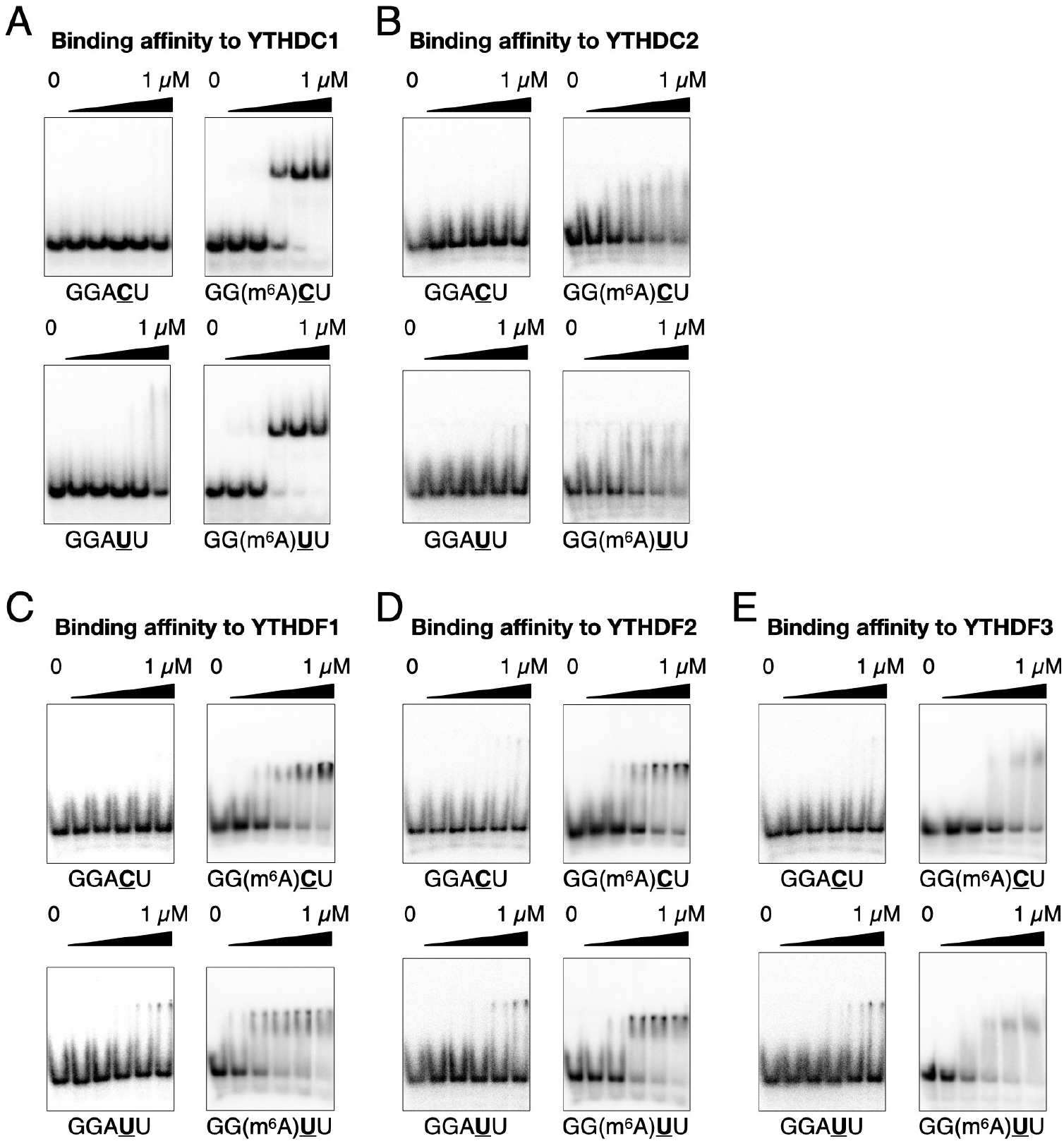
YTH family of m^6^A reader proteins can detect methylated GGAC or GGAU. (**A** to **E**) Electrophoretic Mobility Shift Assay with the YTH domain of YTHDC1 (A), YTHDC2 (B), YTHDF1 (C), YTHDF2 (D), and YTHDF3 (E) and the *MALAT1-derived* RNA fragments containing GGAC or GGAU motifs with or without m^6^A modification. Purified YTH protein concentrations are from left to right, 0 μM, 0.065 μM, 0.130 μM, 0.260 μM, 0.521 μM and 1.04 μM.

**Fig. S6.**
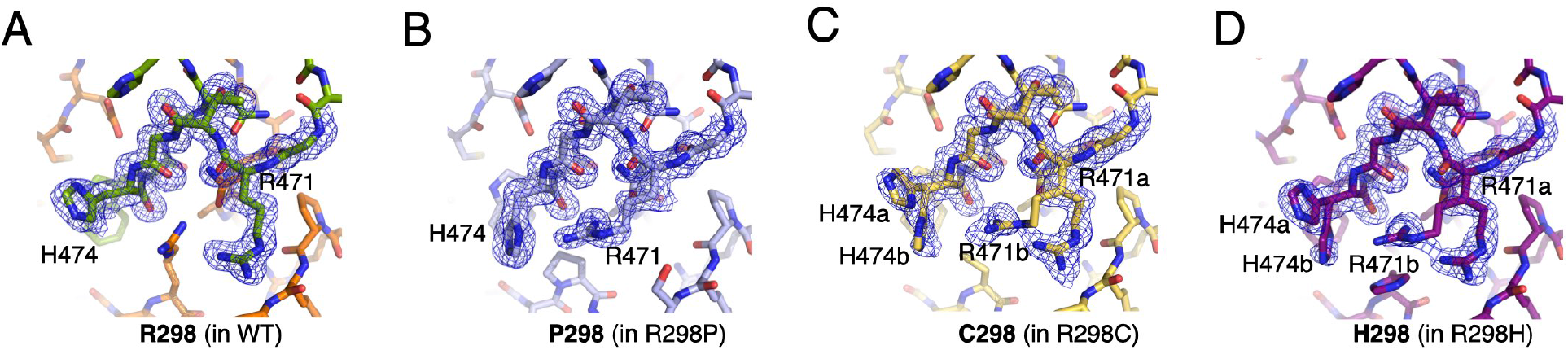
Alternative conformations of nearby side chains with substitutions of METTL14^R298^. (**A** to **D**) WT and mutant structures are individually shown with the respective 2mFo-DFc map contoured at 0.9 σ level. The map is shown only around the residues H474 and R471 of METTL3 for clarity.

**Table S1:** Data collection and structure refinement statistics.

| Data Collection |  |  |  |
| --- | --- | --- | --- |
| Protein | METTL3 and<br>METTL14<sup>R298P</sup> | METTL3 and<br>METTL14<sup>R298H</sup> | METTL3 and<br>METTL14<sup>R298C</sup> |
| Wavelength (Å) | 0.97938 | 0.97938 | 0.97938 |
| Resolution range (Å) | 50-1.65 (1.71-1.65)<sup>a</sup> | 50-1.85 (1.92-1.85) | 50-1.80 (1.86-1.80) |
| Space group | P41212 | P41212 | P41212 |
| Unit cell (Å, °) | 101.4, 101.4, 117.8, 90,<br>90, 90 | 101.4, 101.4, 117.3, 90,<br>90, 90 | 99.7, 99.7, 118.4,<br>90, 90, 90 |
| Unique reflections | 74404 | 52967 | 56123 |
| Multiplicity | 10.7 (9.1) | 15.8 (13.3) | 11.7 (11.2) |
| Completeness (%) | 99.6 (98.9) | 100 (100) | 100 (100) |
| Mean I/sigma (I) | 23.9 (2.5) | 31.4 (2.1) | 23.6 (2.2) |
| Rmerge | 0.097 (0.892) | 0.094 (1.24) | 0.100 (1.08) |
| Structure refinement |  |  |  |
| R-factor/ R-free<sup>b</sup> | 0.1666/ 0.1877 | 0.1711/0.2020 | 0.1928/0.2189 |
| RMS (bonds) | 0.007 | 0.009 | 0.008 |
| RMS (angles) | 0.843 | 0.933 | 1.029 |
| No. of atoms | 4484 | 4387 | 4293 |
| Macromolecules | 4020 | 4005 | 3924 |
| Waters | 464 | 382 | 369 |
| Average B-factor | 24.6 | 29.7 | 25.0 |
| Macromolecules | 23.6 | 29.4 | 27.5 |
| Waters | 32.8 | 30.8 | 28.4 |
| Ramachandran favored (%) | 98.2 | 97.9 | 98.1 |
| Ramachandran outliers (%) | 0 | 0 | 0 |

<sup>a</sup> The values for the data in the highest resolution shell are shown in parentheses.
<sup>b</sup> Rfree = $\sum \text{Test } |F_{\text{obs}}| - |F_{\text{calc}}| / \sum \text{Test } |F_{\text{obs}}|$ , where "Test" is a test set of about 5% of the total reflections randomly chosen and set aside prior to refinement for the structure.

**Table S2:**
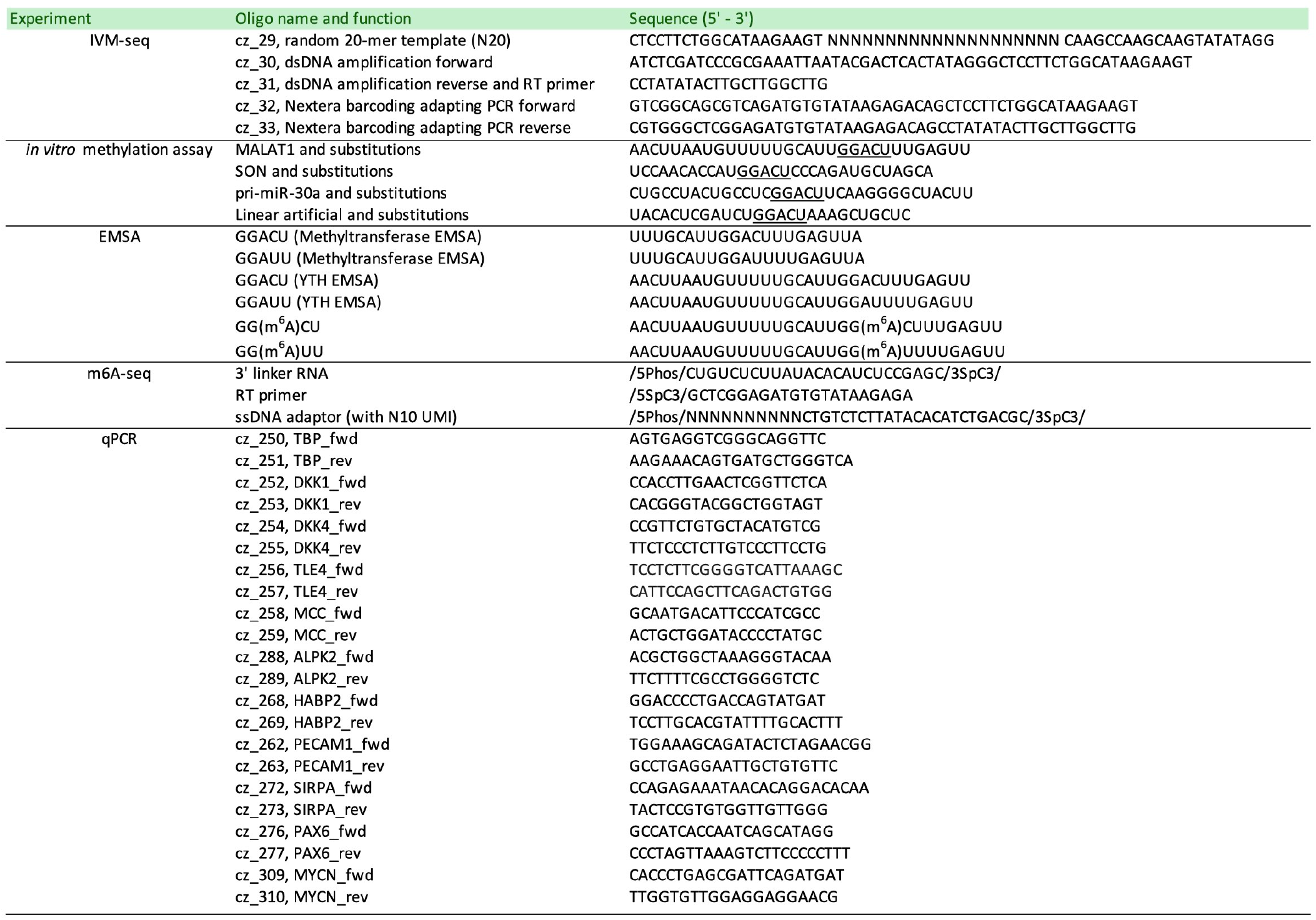
Oligo sequences.

